# Single cell RNA sequencing of blood antigen-presenting cells in severe Covid-19 reveals multi-process defects in antiviral immunity

**DOI:** 10.1101/2020.07.20.212837

**Authors:** Melissa Saichi, Maha Zohra Ladjemi, Sarantis Korniotis, Christophe Rousseau, Zakaria Ait-Hamou, Lucile Massenet, Elise Amblard, Floriane Noel, Yannick Marie, Delphine Bouteiller, Jasna Medvedovic, Frédéric Pène, Vassili Soumelis

## Abstract

COVID-19 can lead to life-threatening acute respiratory failure, characterized by simultaneous increase in inflammatory mediators and viral load. The underlying cellular and molecular mechanisms remain unclear. We performed single-cell RNA-sequencing to establish an exhaustive high-resolution map of blood antigen-presenting cells (APC) in 7 COVID-19 patients with moderate or severe pneumonia, at day-1 and day-4 post-admission, and two healthy donors. We generated a unique dataset of 31,513 high quality APC, including monocytes and rare dendritic cell (DC) subsets. We uncovered multiprocess and previously unrecognized defects in anti-viral immune defense in specific APC compartments from severe patients: i) increase of pro-apoptotic genes exclusively in pDC, which are key effectors of antiviral immunity, ii) sharp decrease of innate sensing receptors, TLR7 and DHX9, in pDC and cDC1, respectively, iii) down-regulation of antiviral effector molecules, including Interferon stimulated genes (ISG) in all monocyte subsets, and iv) decrease of MHC class II-related genes, and MHC class II transactivator (CIITA) activity in cDC2, suggesting a viral inhibition of antigen presentation. These novel mechanisms may explain patient aggravation and suggest strategies to restore defective immune defense.

## Introduction

Severe Acute Respiratory Syndrome-Coronavirus-2 (SARS-Cov-2) infection is at the origin of coronavirus disease 2019 (COVID-19), characterized by a first phase of benign flulike symptoms followed by an efficient control of the infection in most cases. In a second phase, disease aggravation may lead to acute respiratory failure, sepsis, and death ^1–3^. This is due to a multiplicity of factors, which may act separately or in combination: i) exacerbated inflammatory reaction, with both systemic and organ-specific manifestations, ii) persistent viral load, and iii) defective anti-viral defense pathways ^4–6^. Identifying the underlying cellular and molecular mechanisms is of paramount importance to understand COVID-19 physiopathology and guide the development of appropriate therapies.

As of today, studies have characterized the systemic inflammatory response, revealing an excess production of inflammatory cytokines such as IL-6 and IL-1*β* ^6–8^, while the role of TNF-*α* remains less clear^9^. The endothelium may also contribute to the overt inflammatory reaction through the production of soluble mediators^10–11^. Insight also came from targeted immunotherapies, in particular anti-IL-6 compounds, which gave promising results in severe COVID-19^12,13^.However, the cellular source and mechanisms underlying the excessive inflammatory response remain mostly unknown.

Another unresolved question relates to the inefficiency of the innate and adaptive immune system to control the virus in severe COVID-19 patients. It was suggested that production of interferon (IFN)-*α*, a major anti-viral cytokine, was decreased in the most severe patients, as compared to those with moderate disease^8^. However, a recent study argued that an increased IFN-*α* production might contribute to the pathogenic inflammatory response^14^. Other anti-viral mechanisms and their cellular source yet remain to be studied.

Dendritic cells (DC) form a family of innate antigen (Ag)-presenting cells (APC) contributing to the control of pathogens, and subsequent presentation of pathogen-specific Ag to T cells ^15–16^. Their study is challenging for three main reasons: i) they are found in very low numbers in the circulation and in tissue, ii) they lack specific lineage-defining markers, iii) they include an ever-increasing number of subsets^16–17^.All DC subsets may potentially and variably contribute to modulating inflammatory response following viral sensing, producing anti-viral effector molecules, and priming Ag-specific adaptive immune response^17–19^. Plasmacytoid pre-DC (pDC) is a particular subset specialized in anti-viral immunity through the production of large amounts of type I IFN^20–21^. Despite their central role in anti-viral defense, DC contribution to severe COVID-19 pathogenesis is not known.

In this study, we performed a high-resolution single cell RNA sequencing (scRNAseq) analysis of all APC subsets in the peripheral blood of COVID-19 patients. A pre-enrichment step enabled the characterization of even rare DC subsets, not captured in previous scRNAseq studies^14,22^. This revealed previously unrecognized multi-process defects in anti-viral defense.

## Results

### APC subset distribution in COVID-19 patients

To characterize the state and molecular profile of circulating APC, we performed scRNAseq of APC-enriched PBMC from three moderate (non-mechanically ventilated, O2<10L/mn), and four severe (mechanically-ventilated or O2≥10L/mn) COVID-19 patients, at days 1 and 4 post-hospital admission, as well as two healthy controls (HC) (**Fig. 1a and Supplementary Fig. 1a**) (**Table 1 and Supplementary Table 1**). For each sample, 25,000 cells (approximately 20,000 monocytes and 5,000 total DC) were initially loaded to the 10X lane (10X Genomics technology) (**Figure 1a**). We effectively sequenced an average of 9696 cells/sample with a mean value of total genes sequenced per cell estimated at 1638 genes/cell. Samples were split into a discovery dataset (n=8 samples from n=4 patients and n=1 HC) (**Fig. 1-8 and Supplementary Fig. 6**), and a validation dataset (n=6 samples from n=3 patients and n=1 HC) (**Supplementary Fig. 1-5**) (see Material and Methods).

**Table 1.** Patient characteristics at baseline and day 4

| Characteristics | N=7 patients |
| --- | --- |
| Male gender | 6 |
| Age, years | 75 ± 14 |
| Pneumonia severity |  |
| Severe | 4 |
| Mild to moderate | 3 |
| Baseline sample |  |
| Invasive ventilatory support | 3 |
| Concurrent bacterial infection | 0 |
| Total leucocyte count, G/L | 6.62 ± 2.25 |
| Neutrophils, G/L | 5.10 ± 1.80 |
| Monocytes, G/L | 0.87 ± 1.06 |
| Lymphocytes, G/L | 0.87 ± 0.32 |
| Platelet count, G/L | 244 ± 19 |
| C-reactive protein, mg/L | 167 ± 78 |
| Procalcitonin, µg/L | 0.31 ± 0.32 |
| Day 4/5 sample |  |
| Invasive ventilatory support | 3 |
| Concurrent bacterial infection | 2 |
| Total leucocyte count, G/L | 7.42 ± 0.88 |
| Neutrophils, G/L | 5.81 ± 0.77 |
| Monocytes, G/L | 0.68 ± 0.30 |
| Lymphocytes, G/L | 0.99 ± 0.52 |
| Platelet count, G/L | 297 ± 97 |
| C-reactive protein, mg/L | 106 ± 97 |
| Procalcitonin, µg/L | 1.86 ± 2.88 |

**Figure 1.**
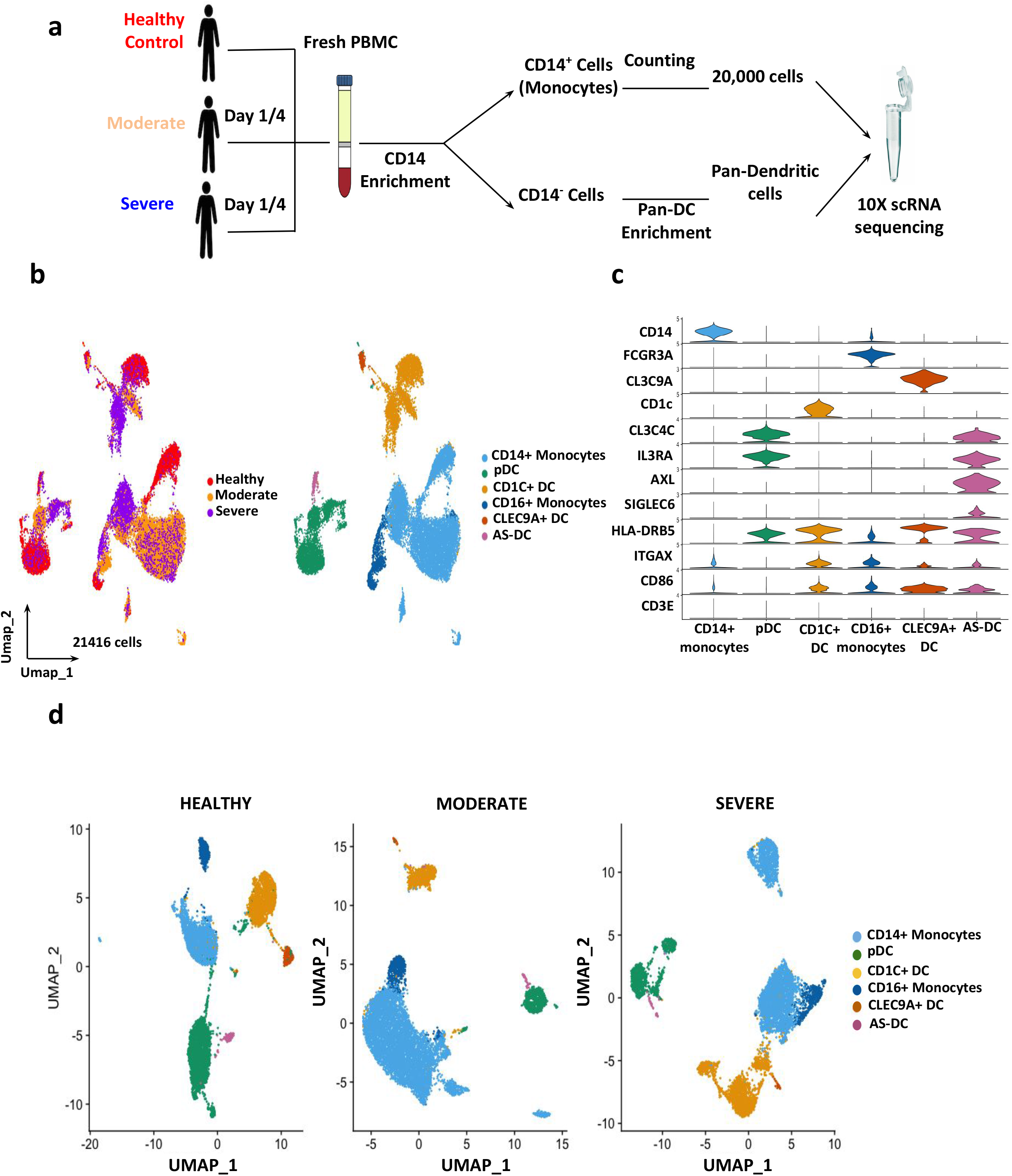
Circulating APC subset diversity in COVID-19. **a.** Schematic representation of the experimental workflow. APC were enriched from fresh PBMC of healthy donors and COVID-19 patients, with either moderate or severe clinical symptoms at both day 1 and day 4 post-hospital admissions. 25,000 total APC per sample were sequenced using 10X Genomics, **b**. Cellular map of APC subsets at the single-cell resolution level displayed on UMAP (Uniform Manifold Approximation and Projection) dimension reduction based either on severity (left) or identified cell types (right). Analysis was performed on all samples from the discovery dataset, **c**. Identification of APC subsets defined by gene expression of standard subset-specific markers, **d**. UMAP plot of detected APC populations for each severity group (Healthy controls, moderate and severe patients).

Uniform manifold approximation and projection (UMAP) and graph-based clustering of all severity group samples allowed us to manually annotate clusters with their cellular identities (**Fig. 1b**). Using dimensionality reduction, we established a map of 21,416 APC (discovery set) identifying 6 subsets: 11,805 CD14^+^ and 1,354 CD16^+^ monocytes, 3,958 pDC, 301 CLEC9A^+^ and 3,785 CD1c^+^ DC (cDC1 and cDC2, respectively), and 213 Axl-Siglec 6 (AS)-DC (**Fig. 1b and Supplementary Fig. 1b**). These APC populations were captured across the whole dataset (**Supplementary Fig. 1a)**. Their identity was confirmed by the expression of unique combinations of markers (**Fig. 1c**). CD14^+^ monocytes expressed exclusively their lineage-defining CD14 marker, whereas CD16^+^ monocytes expressed FCGR3A. As expected, all DC populations expressed higher levels of HLA-DR and CD86, as compared to monocytes (**Fig. 1c**). AXL expression distinguished AS-DC from pDC, whereas CD1c and CLEC9A characterized the respective cDC subsets. The absence of CD3 *ε* confirmed the lack of T cell contamination (**Figure 1c**). Dimensionality reduction and graph-based sub-clustering allowed us to retrieve all APC subsets across patient groups. Proportions, however, were different, with a relative decrease in pDC, and increase in CD14^+^ monocytes, in COVID-19 patients as compared to HC (**Fig. 1d and Supplementary Fig. 1c**). Interestingly, we denoted an increased heterogeneity among pDC, CD14^+^ monocytes, and CD1C^+^ DC in severe patients (**Fig. 1d**). Overall, our enrichment strategy allowed the efficient identification of all APC populations including the rare pDC, AS-DC, and CLEC9A^+^ DC, enabling further molecular and phenotypic characterization.

### Inflammatory pathways in COVID-19 APC

We used unsupervised pathway enrichment methods on total APC, in order to identify functional pathways associated with COVID-19-related inflammation. We detected 743 differentially expressed genes (DEG) between HC, moderate and severe COVID-19 patients, with an absolute fold change higher than 1.4 (**Fig. 2a**). Among DEG, 184 genes were up-regulated in HC, 158 in moderate, and 16 in severe, as compared to the two other groups, respectively (**Fig. 2b).** We performed pathway enrichment analysis on DEG using hallmark signatures compiled in MsigDB^23^, and focused on pathways enriched in each patient group, as compared to HC. “KRAS signaling”, previously associated with down-regulation of IFN-induced anti-viral response ^24^, and “cholesterol homeostasis” pathways, were enriched in moderate and severe patients (**Fig. 2c**). Both moderate and severe patients displayed enriched “Hypoxia”, “Apoptosis”, “Inflammatory response”, and “TNF-*α* signaling” pathways (**Fig. 2c**). Differential expression of those pathways was retrieved in our validation set (**Supplementary Fig. 2**). Pathway-level analyses were supported by the coordinated up-regulation of many genes involved in TNF-*α* and Hypoxia signaling (**Fig. 2d and e**).

**Figure 2.**
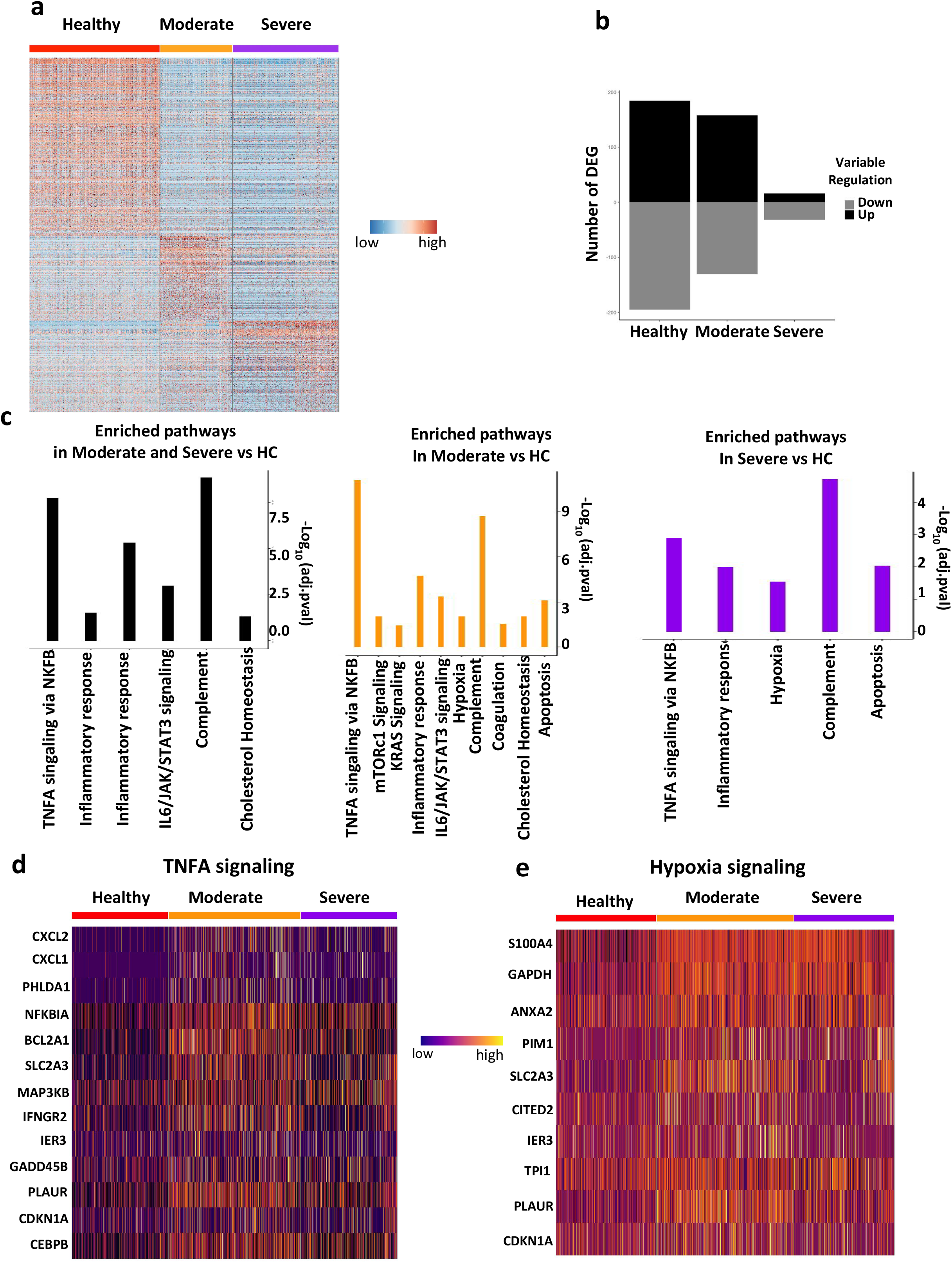
Global increase in inflammation-associated pathways in COVID-19 APC. **a**. Heatmap representation of differentially expressed genes (DEG) between severity groups. Healthy control (n sample=1), moderate (n sample =3) and severe (n sample =4) (discovery dataset), **b**. Histogram representation of DEG between severity groups. Up-regulated genes are shown in black and down-regulated genes in grey, **c**. Gene set enrichment analysis of pathways up regulated in: moderate and severe patients compared to HC (left), moderate compared to HC (middle), and severe compared to HC (right). Pathway enrichment analysis is shown in Material and Methods, **d-e**. Heatmap representation of genes differentially expressed contributing to either **d**. TNFA signaling or **e**. Hypoxia signaling pathways

### Defective inflammatory cytokine and IFN responses in COVID-19 APC

Next, we addressed the global contribution of APC to the expression of specific inflammatory cytokines and their receptors known to be involved in COVID-19 physiopathology. As compared to HC, the pro-inflammatory genes CXCL8 and IL1B increased in both severity groups, whereas CCL3 and CXCL2 were up-regulated in moderate patients (**Fig. 3a and Supplementary Fig. 3a**). In parallel, we detected a decrease of TGFB1 and IL18, in both moderate and severe patients, together with a decrease in the expression of receptors for IL-6, IL-10 and TNF-*α* (**Fig. 3a and Supplementary Fig. 3a**). Next, we explored biological pathways associated to downstream cytokine signaling. In both moderate and severe patients, we detected higher scores for “p53”, “inflammatory response”, “TGF-*β* signaling”, and “IL-6/JAK/STAT3 signaling” pathways, as compared to HC (**Fig. 3b**).

**Figure 3.**
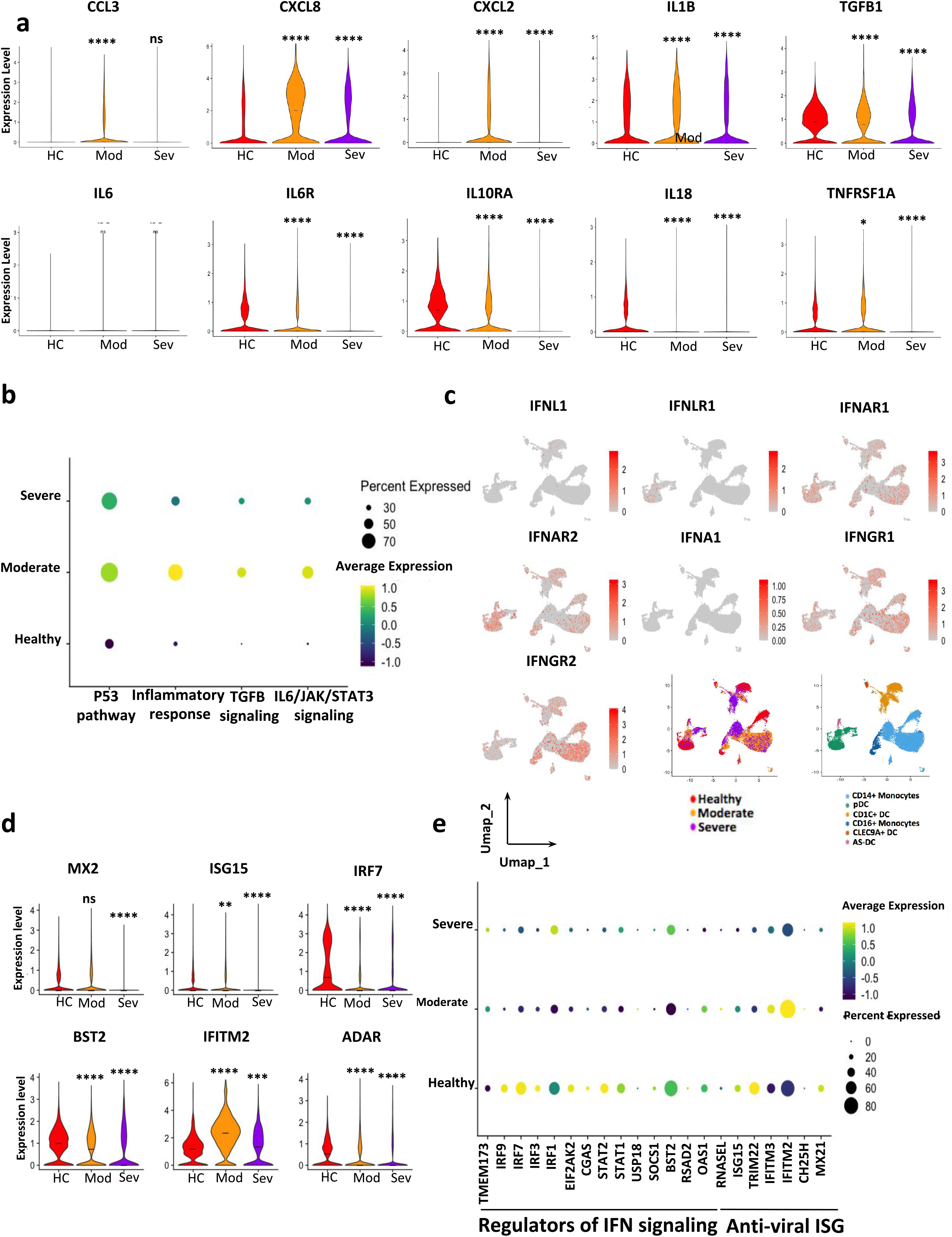
Defective cytokine and IFN responses in severe COVID-19 APC. **a.** Violin plot representation of gene expression for cytokines (IL1B, TGFB1, IL-6 and IL-18), chemokines (CCL3, CXCL8, CXCL2) and receptors (IL6R, IL10R and TNFRSF1A) detected by scRNAseq and comparison between severity groups, **b**. Dot plot depicting the enrichment scores of pathways downstream of IL-1B, IL-6 and TGFB1 inflammatory cytokines. Score levels are color-coded; the percentage of cells expressing the respective score pathway is size coded, **c.** Feature plot of IFNs and their receptor expression across all cell types and severity cases within UMAP (discovery dataset). Expression levels are color-coded, **d.** Violin plot representation of anti-viral IFN stimulating genes between severity groups. **e.** Dot plots of regulators of IFN signaling and anti-viral IFN stimulating genes in healthy control, moderate and severe patients. Expression levels are color-coded; Percentage of cells expressing the respective gene is size coded. Asterisks above severe indicate *P* values for severe versus control; asterisks above moderate indicate significance of moderate versus control. \**P* < 0.05, \*\**P* < 0.01, \*\*\**P* < 0.001.

IFN family of cytokines is one of the most important mediators of anti-viral responses. Using previously shown UMAP coordinates, we represented the expression level of IFNL1, IFNL1R, IFNAR1, IFNAR2, IFNA1, IFNGR1 and IFNGR2, according to the severity and cell type (**Fig. 3c**). IFNL1 and IFNL1R were marginally detected in our discovery dataset (**Fig. 3c**), while absent from the validation dataset (**Supplementary Fig. 3b**), supporting that IFN-*λ* pathway was not significantly expressed. IFNAR1 and IFNAR2 were detected in almost all identified cell populations, with higher levels in pDC from HC, and CD14^+^ monocytes from both HC and moderate patients (**Fig. 3c and Supplementary Fig. 3b**). Severe patients expressed much lower levels of both IFNAR1 and 2, suggesting a defect in IFN-*α* signaling (**Fig. 3c**). IFNGR1 and IFNGR2 were highly expressed by CD14^+^ monocytes and CD1c^+^ DC, and lower in pDC from HC and moderate patients (**Fig. 3c**). Type I IFN, such as IFNA1, were not detected in any of the APC subsets and patient groups (**Fig. 3c and Supplementary Fig. 3b**).

To further investigate the involvement of IFN-stimulated genes (ISG), we quantified the expression level of six major ISG (MX2, ISG15, IRF7, BST2, IFITM2 and ADAR) ^25^. Severe patients displayed low levels of all these genes, suggesting that the development of severe COVID-19 might be related to IFN dysregulation (**Fig. 3d and Supplementary Fig. 3c**). ISG have been separated into “anti-viral ISG” and “Regulators of IFN signaling”^25^ . Both moderate and severe patients APC expressed lower levels of most genes in these two categories, suggesting a global perturbation of IFN downstream functions (**Fig. 3e**).

### Increased apoptosis and multi-process effector defects in severe COVID-19 pDC

After having analyzed perturbations in COVID-19 APC at the global level, we sought to decipher alterations occurring in specific APC subsets. Sub-clustering of the pDC subset identified distinct pDC clusters for each severity group (**Fig. 4a**). DEG among pDC from the three patient groups revealed signatures associated to HC versus COVID-19 patients **(Fig. 4b)**. Pathway enrichment analysis revealed an enrichment of “Hypoxia” and “TNF-*α* signaling via Nuclear Factor (NF) kB”, in pDCs from moderate and severe, as compared to HC (**Fig. 4c**). PDC from severe patients exhibited specific enrichment in a number of pathways, including “apoptosis”, “p53 pathway”, “IL6/JAK/STAT3 signaling”, and “IL2/STAT5 signaling” (**Fig. 4c**). To accurately compare differentially enriched pathways among severity groups, we computed pathway-specific module scores using the gene lists corresponding to each hallmark signature. This confirmed that pDC from severe patients displayed significantly elevated scores for “Pro-apoptosis”, “IFNG response”, “Hypoxia”, and “TNFa signaling” pathways, as compared to HC (**Fig. 4d**). Overall, these analyses converge to establish a global deficiency in pDC through induced apoptosis, as well as a pro-inflammatory state related to cytokine-induced NFkB and JAK/STAT signaling.

**Figure 4.**
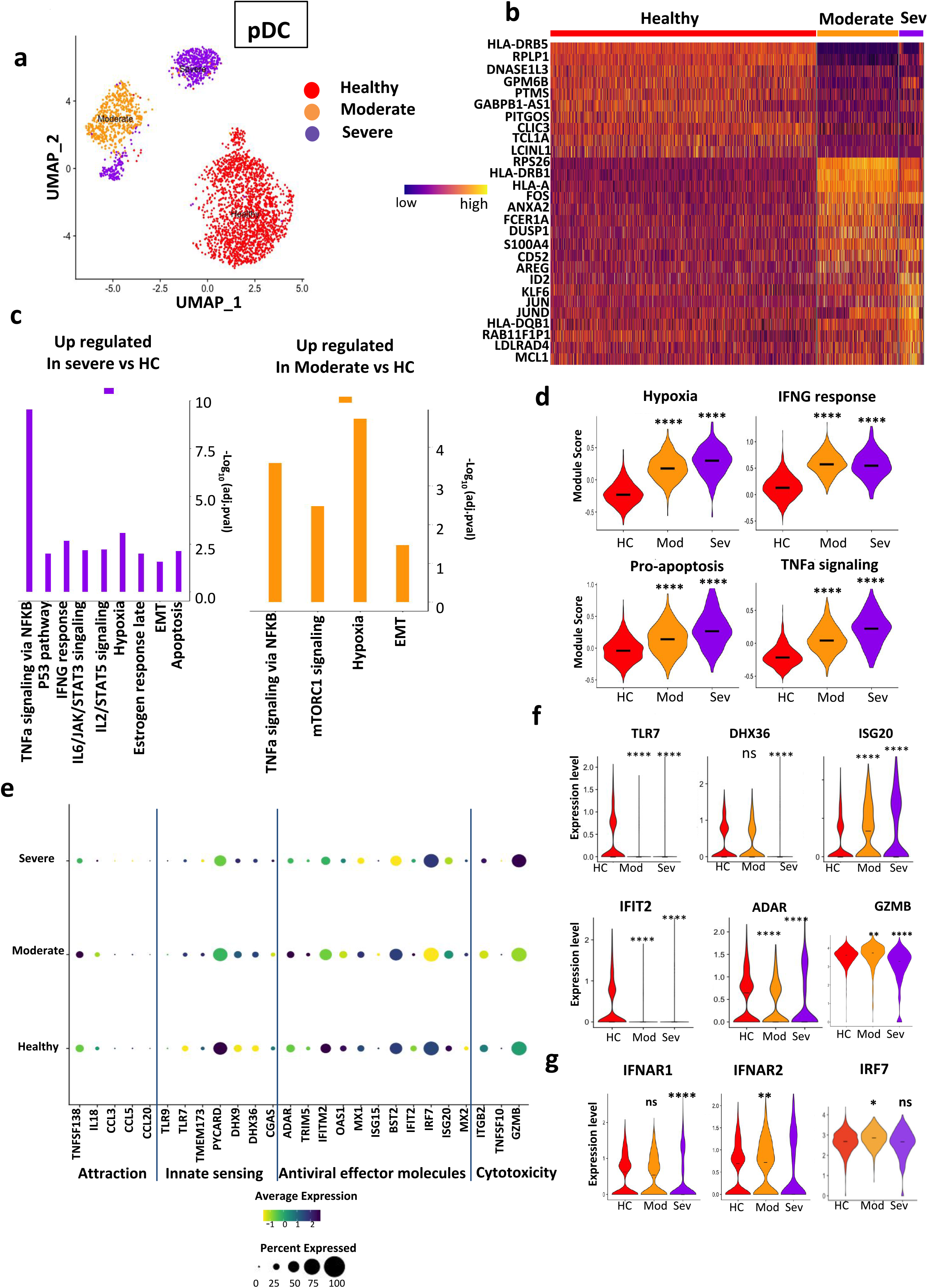
Multiprocess defects in severe COVID-19 pDC effector pathways. **a**. UMAP displaying identified pDC in all severity cases (discovery dataset), **b**. Heatmap depicting differentially expressed genes between identified pDC derived from HC, moderate and severe patients, **c**. Gene set enrichment analysis of pathways up-regulated in severe versus HC (purple), moderate versus HC (orange). Pathway enrichment analysis is described in Material and Methods, **d**. Violin plot representation of signature scores for “Hypoxia”, “IFNG response”, “pro-apoptosis” and “TNFA signaling” pathways, between severity groups, **e**. Dot plot of in-house constructed pDC-related functional modules (cytotoxicity, anti-viral effector molecules, inate sensing, and attraction) and comparison between severity groups, Expression levels are color-coded; Percentage of cells expressing the respective gene is size-coded, **f**. Violin plot representation of genes involved in anti-viral immune responses by pDC between severity groups. Asterisks above severe indicate *P* values for severe versus control; asterisks above moderate indicate significance of moderate versus control. \**P* < 0.05, \*\**P* < 0.01, \*\*\**P* < 0.001.

We asked whether apoptosis and pro-inflammatory signaling signatures altered pDC effector functions. We defined four modules including immune effector molecules, using a literature-driven manual curation: immune cell attraction (hereafter “attraction”) (18 genes), “innate sensing” (12 genes), “anti-viral effector molecules” (23 genes), and “cytotoxicity” (12 genes) (**Fig. 4e**). Each of these modules was crossed with the pDC expression matrix, and detected genes were depicted for each patient group based on average expression and percentage of positive cells (**Fig. 4e**). Only five genes were detected as part of the “attraction” module, with minor differences between groups, showing that this function is not significantly represented in HC, and COVID-19 pDC at this advanced-stage of the disease. On the contrary, many genes in the “innate sensing”, “anti-viral effector molecules”, and “cytotoxicity” modules were detected in the three groups, and followed the same pattern: baseline levels in HC, increased levels in moderate, and down-regulation in severe (**Fig. 4e and f**) (**Supplementary Fig. 4a)**. This was particularly striking for the viral sensors TLR7, DHX9, DHX36, and the cytotoxic molecule GZMB.

Last, we sought to identify potential mechanisms underlying the defective pDC functions in severe patients. Transcription factors controlling IFN-*α* production may be involved, among which IRF-7^20^. Although we found that IRF7 in moderate patients pDC was increased as compared to HC, expression was back to baseline in severe patients pDC (**Fig. 4g**). We turned to IFNAR expression as an alternative hypothesis, since IFNAR signaling is key to drive downstream anti-viral IFN-induced molecules. We found that IFNAR1 and 2 were significantly decreased in pDC from severe patients, as compared to those from moderate patients and HC (**Fig. 4g**) (**Supplementary Fig. 4b)**.

### Coordinated transcriptional adaptation in COVID-19 monocyte subsets

Monocytes are important inflammatory cells and have been implicated in the physiopathology of severe sepsis and COVID-19. In order to get insight into monocyte diversity, we used UMAP to visualize total monocytes according to disease severity (**Fig. 5a, left plot**), and subset diversity (**Fig. 5a, right plot**). We observed that CD14^+^ and CD16^+^ monocytes were clearly distinct in HC, but more similar in patients (**Fig. 5a).** This suggested that COVID-19 might imprint common transcriptional programs to both subsets. To investigate alterations specifically encountered in CD14^+^ monocytes, we performed dimensionality reduction through Independent Component Analysis (ICA), and highlighted cells according to their severity group. We observed that IC1 clearly separated HC from COVID-19 monocytes, whereas IC3 distinguished severe CD14^+^ monocytes from the remaining cells (**Fig. 5b**).

**Figure 5.**
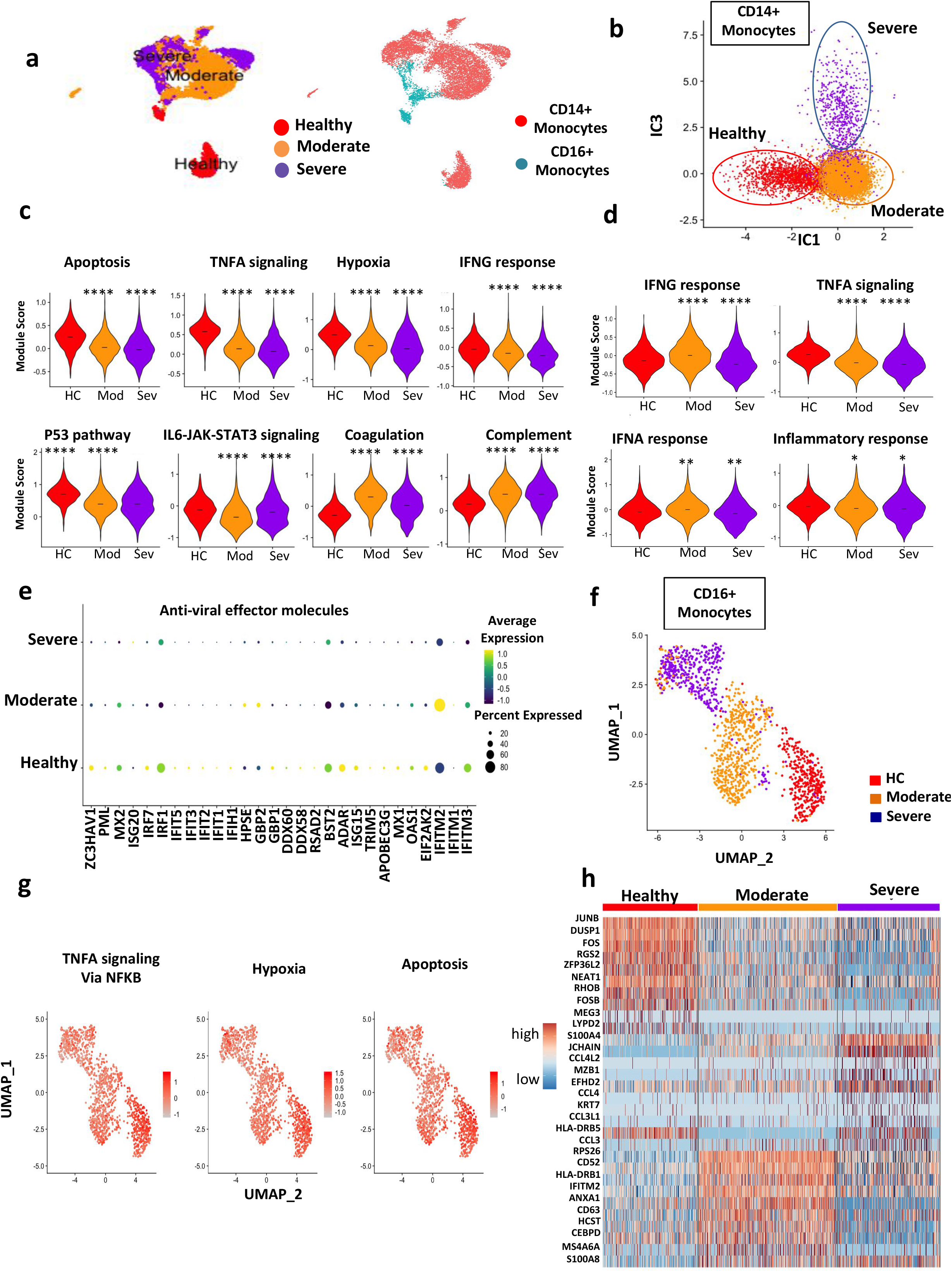
Dissection of inflammatory and anti-viral response pathways in monocyte subsets. **a**. UMAP displaying the detected monocyte subsets (CD14^+^ and CD16^+^) for all patient groups in the discovery dataset, **b**. Independent Component Analysis (ICA) representation of CD14^+^ monocytes derived from all severity groups. IC1 and IC3 components allowed the separation of CD14^+^ monocytes according to severity, **c**. Violin plot for the expression of enriched pathways in top50 genes characterizing IC1, **d**. Violin plot for the expression of enriched pathways in top50 genes characterizing IC3, **e.** Dot plot of in-house constructed anti-viral effector molecule modules across CD14^+^ monocytes and severity groups. Expression levels are color-coded; Percentage of cells expressing the respective gene is size coded; **f.** UMAP displaying CD16^+^ monocytes derived from all severity groups, **g.** UMAP representation of pathways score enriched for TNFA signaling, Hypoxia and Apoptosis. The score level is color-coded and described in Material and Methods, **h.** Heatmap representation of top 10 DEG between severity groups. Asterisks above severe indicate *P* values for severe versus control; asterisks above moderate indicate significance of moderate versus control. \**P* < 0.05, \*\**P* < 0.01, \*\*\**P* < 0.001.

Pathway enrichment analysis on the top50 genes contributing positively and negatively to IC1, followed by estimation of a score for each functional module, identified key pathways that segregated COVID-19 CD14^+^ monocytes from HC (**Fig. 5c**). Among them, we found decreased “apoptosis” and “p53 signaling”, decreased “hypoxia”, “TNFA signaling”, and “IL6-JAK-STAT3 signaling », whereas « complement » and « coagulation » pathways were increased, suggesting a possible contribution to vascular events in the physiopathology of the disease (**Fig. 5c**). Similar analyses were done on IC3 top50 genes, and revealed lower “IFNA response” and “inflammatory response” in severe patients (**Fig. 5d**). Last, we asked whether decreased apoptosis in severe CD14^+^ monocytes would be associated to sustained or increased antiviral effector functions. We used our manually curated gene functional module, and estimated expression levels across patient groups (**Fig. 5e**). We observed a global decrease of almost all anti-viral effector molecules in severe patients as compared to HC (**Fig. 5e**) or moderate patients (**Supplementary Fig. 5a**). Overall, CD14^+^ monocytes showed defective anti-viral functions, while possibly contributing to vascular inflammation by triggering coagulation events.

In parallel, we sub-clustered CD16^+^ monocytes and reduced the data dimension using UMAP projection to depict the corresponding clusters for each severity group (**Fig. 5f**). Pathway enrichment analysis on top50 contributing genes to Umap2 dimension highlighted significantly lower enrichment scores corresponding to “TNFA signaling”, “Hypoxia” and “Apoptosis” pathways in moderate and severe patients, versus HC (**Fig. 5g**). As a complementary approach, we performed differential expression analysis, and identified signatures specific to each of the CD16^+^ monocyte cluster (**Fig. 5h, top 10 DEG for each cluster**). These disease-associated changes in CD16^+^ monocytes paralleled the ones observed in CD14^+^, suggesting common adaptation mechanisms.

### CLEC9A^+^ DC and AS-DC-specific transcriptional alterations

Thanks to our APC-enrichment protocol, we could efficiently recover rare CLEC9A^+^ DC and AS-DC subsets (**Fig. 6**). Sub-clustering of CLEC9A^+^ DC and AS-DC identified clusters specific to HC, moderate, and severe patients (**Fig. 6a and e**), although less clearly defined than pDC, and monocyte sub-clusters (**Fig. 4, 5**). In addition, we identified a strong decrease in CLEC9A^+^ DC proportions in moderate and severe patients (**Supplementary Fig. 1b**).

**Figure 6.**
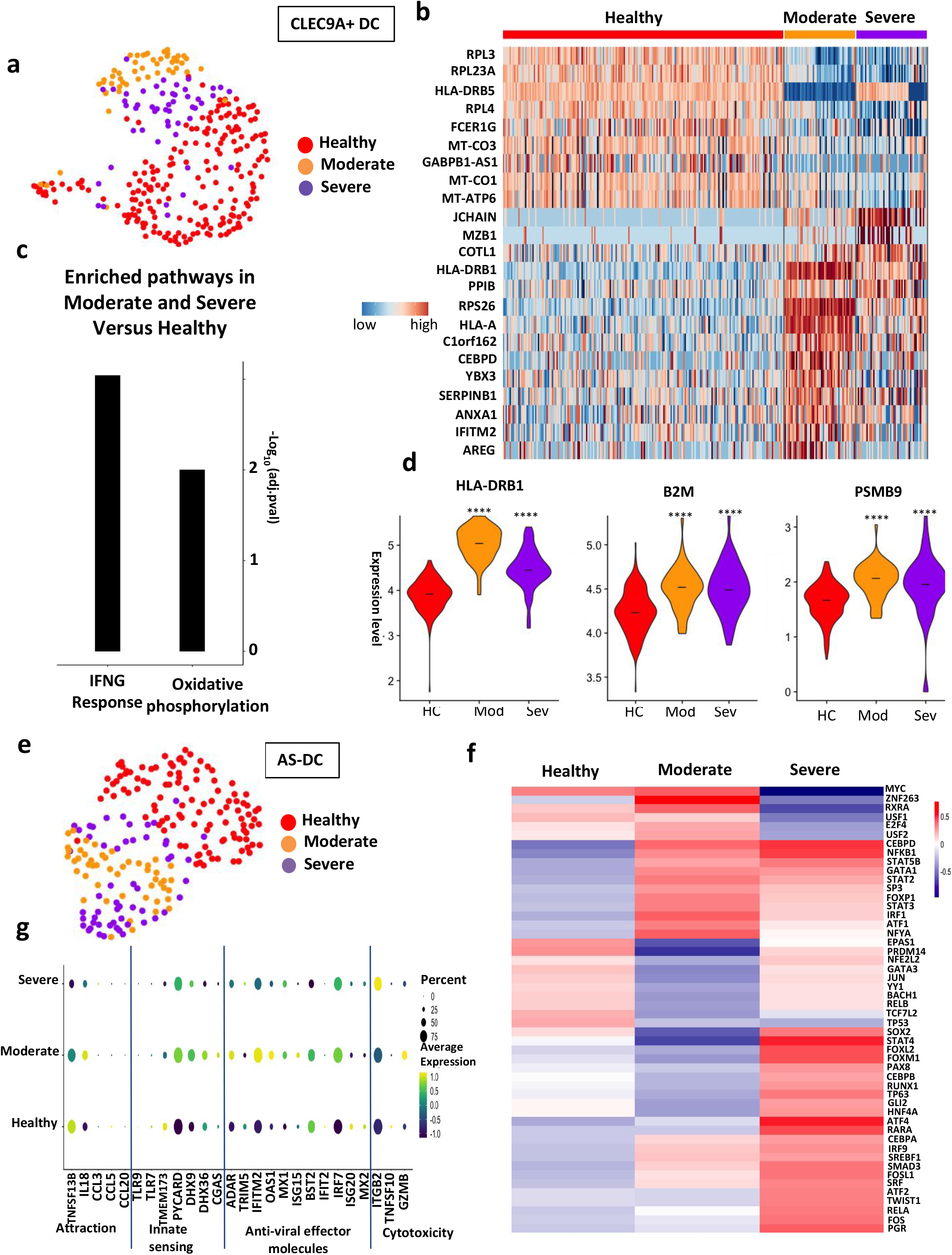
Molecular and functional modules in rare CLEC9A^+^ and AS-DC subsets. **a**. UMAP displaying the CLEC9A^+^ DC subsets of all patients, **b.** Heatmap representation of top 10 DEG between severity groups for CLEC9A^+^ DC, **c.** Histogram representation of enriched pathways derived from up-regulated genes in moderate and severe patients compared to HC (–Log10 (adjusted p value) is represented on the Y axis), **d.** Violin plot representation of representative DEG across CLEC9A^+^ DC clusters, **e.** UMAP displaying the AS-DC subset of all patients, **f.** Heatmap representation of the activity of transcription factors using Dorothea inference tool (Z-scores are represented), **g.** Dot plot of in-house constructed functions (“cytotoxicity”, “anti-viral effector molecules”, “innate sensing” and “attraction”) for AS-DC subset between severity groups; Expression levels are color-coded; Percentage of cells expressing the respective gene is size coded. Asterisks above severe indicate *P* values for severe versus control; asterisks above moderate indicate significance of moderate versus control. \**P* < 0.05, \*\**P* < 0.01, \*\*\**P* < 0.001.

DEG among CLEC9A^+^ DC sub-clusters included specific transcriptional signatures segregating moderate and severe patients from HC (**Fig. 6b**). Up-regulated genes in moderate and severe patients compared to HC showed enrichment in “IFNG” and “Oxidative phosphorylation” pathways (**Fig. 6c**). Specifically, important genes in the IFNG pathway (HLA-DRB1, B2M, PSMB9) were significantly increased in moderate and severe CLEC9A^+^ DC, versus HC (**Fig. 6d**).

In order to characterize molecular pathways in AS-DC sub-clusters, we used two complementary approaches. First, we inferred transcription factor (TF) activity using Dorothea algorithm ^26^ and scored the activity of each regulon using Viper inference tool ^27^. This identified a large number of highly variant TF activity scores, including a group with increased activity in severe patients (**Fig. 6f**). Interestingly, these included STAT4 and RUNX1, which are usually associated to Th1 responses, and may control pro-inflammatory genes in AS-DC. Second, we used our manually curated functional modules to infer functional defects in COVID-19 AS-DC (**Fig. 6g**). As for pDC, the “Attraction” module was not well represented, although two of its members (IL18, TNFSF13B) were decreased in severe patients (**Fig. 6g**). The most important changes were observed for “anti-viral effector molecules”, which increased in moderate patients as compared to HC, and further decreased in severe (**Fig. 6g**), compatible with a compromised anti-viral immune control.

### Downregulation of MHC-II and loss of CIITA activity in CD1C^+^ DC

We next focused on disease-induced alterations in CD1c^+^ DC. Sub-clustering of CD1c^+^ DC identified well-defined clusters specific to HC, moderate and severe patients (**Fig. 7a**). Computation of DEG across the three clusters revealed specific transcriptional signatures (**Fig. 7b**). This included a significant down-regulation of the MHC-II genes HLA-DPB1, HLA-DRB5, and HLA-DQA2, in CD1c^+^ DCs from moderate and severe COVID-19 patients, as compared to HC (**Fig. 7b and c**). HLA-DQA1 and DQB1 were not differentially expressed (not shown). Interestingly, HLA-DRB1 was decreased in severe patients as compared to moderate (**Fig. 7c and Supplementary Fig. 5b**).

**Figure 7.**
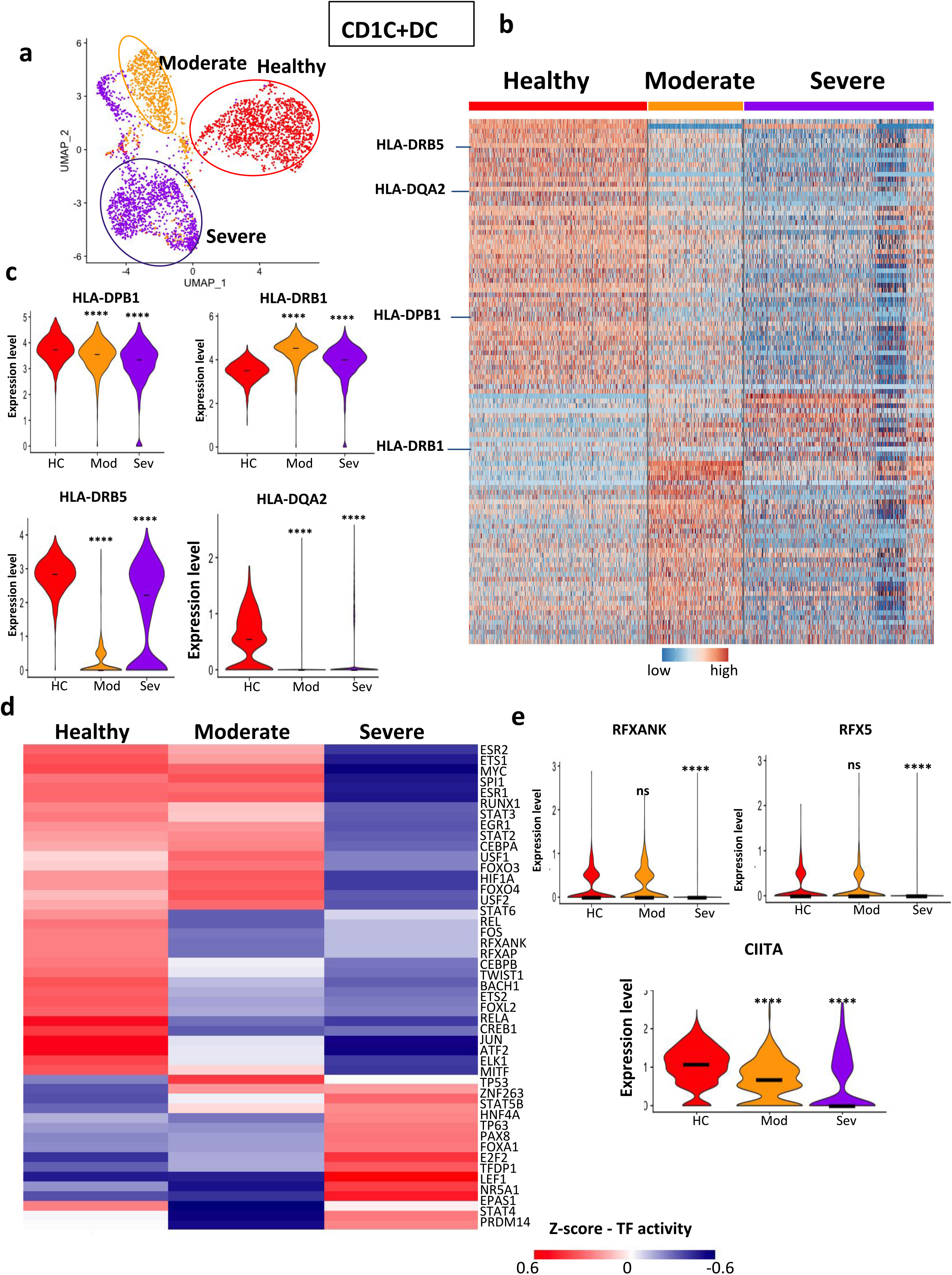
Downregulation of MHC II and upstream transcriptional regulators in severe COVID-19 CD1c^+^ DC. **a.** UMAP displaying CD1c^+^ DC subsets of all patients, **b.** Heatmap representation of DEG between severity groups for CD1c^+^ DC. Highlight of MHC-II-related genes up-regulated in HC compared to COVID-19 patients, **c.** Violin plot representation of MHC-II-related gene expression between severity groups, **d.** Heatmap representation of the activity of transcription factors inferred by Dorothea tool (Z-scores are represented), **e.** Violin plot representation of gene expression of transcription factor regulators of MHC-II-related genes, comparison between severity groups. Asterisks above severe indicate *P* values for severe versus control; asterisks above moderate indicate significance of moderate versus control. \**P* < 0.05, \*\**P* < 0.01, \*\*\**P* < 0.001.

In order to identify upstream mechanisms, we studied the differential estimated activity of TF (**Fig. 7d**), similar to the strategy used for AS-DC (**Fig. 6g**). Three different estimated activity patterns were obtained according to disease severity. First, we found TF activities that were down-regulated in CD1c^+^ DC from moderate and severe patients including REL, recently shown to be a myeloid checkpoint^28^, FOS and JUN. Second, we found TF activities down-regulated in severe patients as compared to HC and moderate patients, including STAT2, 3 and 6, involved in immune responses by controlling transcription of cytokine-induced genes. Third, we identified TF activities up-regulated in severe patients as compared to HC and moderate patients, including E2F2 and TFDP1 that regulate cell cycle and proliferation (**Fig. 7d**). Overall, CD1c^+^ DC from severe COVID-19 patients were characterized by a defect in TF activity involved in cytokine-induced gene expression, and an increased expression of cell cycle regulators.

In order to explain the down-regulation of MHC class II genes, we focused on their upstream TF regulators. We show a significant down-regulation of RFX5 in CD1c^+^ DC from severe patients (**Fig. 7e and Supplementary Fig. 5c**). We also observed a down-regulation of CIITA, and most pronounced in severe patients (**Fig. 7e and Supplementary Fig. 5c**), which may be due to the down-regulation of USF1 (**Fig. 7d)**, interacting with STAT1 to activate IFN-*γ*-induced transcription of the CIITA gene ^29^.

Since MHC-II genes are involved in DC-T cell interaction, we hypothesized that a more global dysfunction of DC-T cell communication may occur in COVID-19 APC. To test this, we applied our cell communication inference computational framework ICELLNET^30^(**Supplementary Fig. 6)**. Using our “reference partner cell” methodology, we inferred potential communication between each of the APC subsets and CD4 T cells, in each of the disease groups (**Supplementary Fig. 6a).** Cell connectivity networks revealed a global decrease in APC-T cell communication in severe, as compared to moderate and healthy, predominantly in CD1c^+^ DC and CD14^+^ monocytes (**Supplementary Fig. 6a and b**). We then explored the various communication molecular families that may explain this decrease. This revealed a dominant contribution of checkpoint molecules and cytokines for CD1c^+^ DC-T cell communication, while cytokines were mostly underlying the decrease in CD14^+^ monocytes-T cell communication (**Supplementary Fig. 6b**). We could further identify ligand-receptor pairs within each of these molecular families, to explain the decreased CD1c^+^ DC-T cell communication: MHC II-LAG3, CD80-CD28, and CD48-CD2 interactions (**Supplementary Fig. 6c**). Among cytokines, IL-10 and TGFB-mediated interactions were predominantly impacted in severe patients, which may contribute to immunopathology through excessive Th1 responses (**Supplementary Fig. 6c**).

### Persistence of multi-process defects in severe COVID-19 APC subsets

Last, we asked whether functional pathway alterations observed across APC subtypes were sustained over time during the course of SARS-CoV-2 infection. We compared scRNAseq datasets generated at day 1 versus day 4 post-hospital admission (**Fig. 8**). In a first approach, unsupervised differential expression of severe APC mean expression genes, between Day 1 and Day 4, revealed a low number of DEG, most of them being mitochondrial genes, not related to any specific functional module (**Fig. 8a**). We hence used an alternative hypothesis-driven approach, by selecting genes involved in previously identified altered functions, in severe COVID-19 APC, and systematically compared Day 1 and Day 4 expression levels, and percent cell expression (**Fig. 8b**). Most Day 1 defects were sustained at Day 4, in particular the low expression of anti-viral effector molecules in pDC and monocytes, as well as the increased apoptosis pathway in pDC (**Fig. 8b**). Interestingly, we noted an increase in two MHC-II molecules, HLA-DQA2 and DRB5, in CD1c^+^ and CLEC9A^+^ DC, between Days 1 and 4 (**Fig. 8b**). This suggests a possible recovery of Ag presentation with severe COVID-19 evolution.

**Figure 8.**
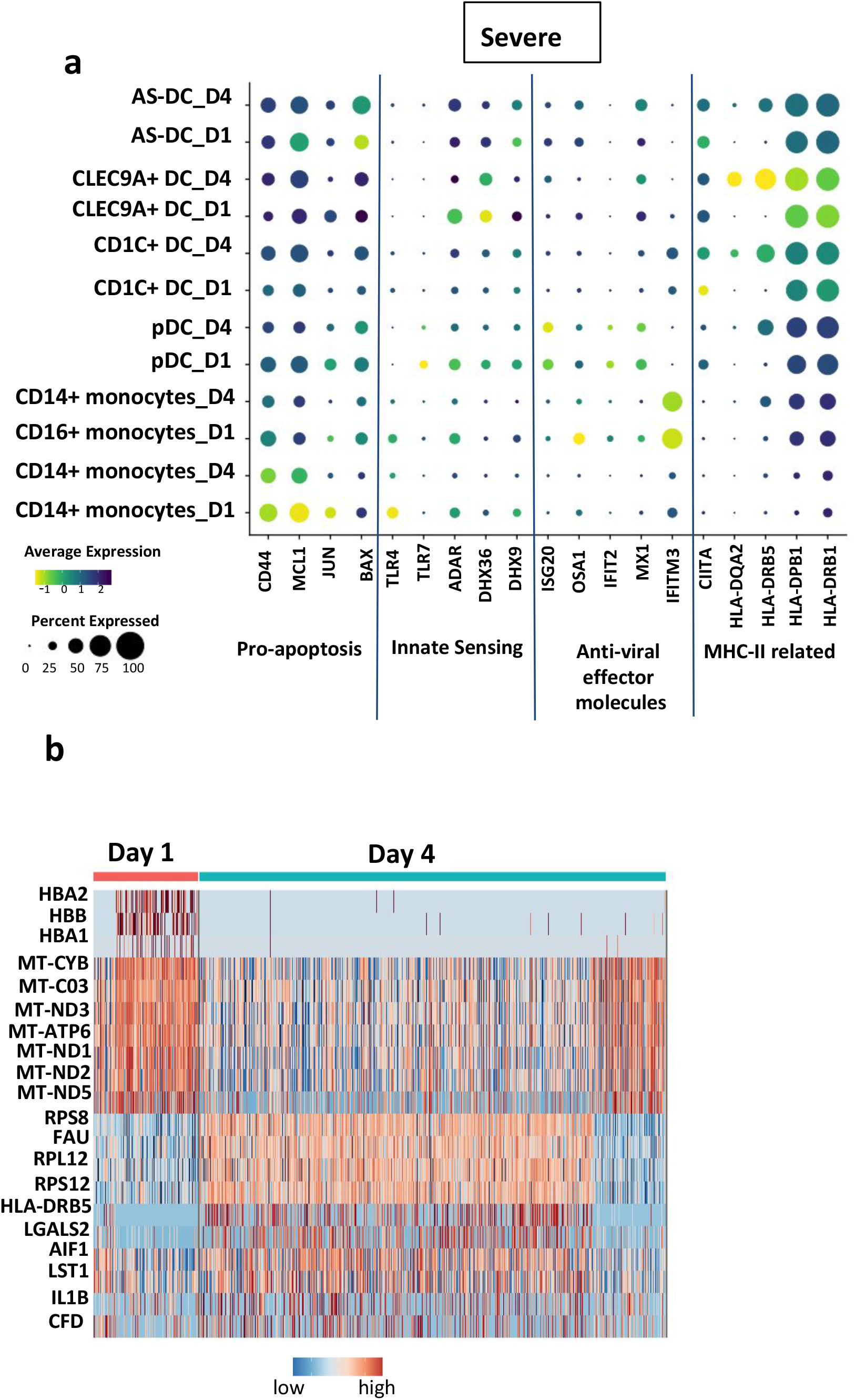
Persistent defects in anti-viral defense pathways in severe COVID-19 APC. **a**. Heatmap representation of top 20 DEG between day-1 and day-4 groups for APC subsets from severe patients, **b.** Dot plot representation of temporal evolution (Day-1, Day-4) of gene expression levels involved in previously identified perturbed pathways (MHC-II related genes, antiviral effector molecules, innate sensing, apoptosis) in specific APC subsets from severe patients. Expression levels are color-coded; Percentage of cells expressing the respective gene is size coded;

## Discussion

Severe COVID-19 harbors a complex physiopathology that only starts to be deciphered. This complexity comes from host-pathogen interactions evolving over time, and involving a large number of underlying cellular and molecular mechanisms. Hence, detailed studies on various immune cell compartments are required to get a global view of the process. DC are central to immune responses by linking innate and adaptive immunity, in particular during infection ^16^. DC are very rare cell types composed of multiple subsets ^17^, justifying dedicated studies to uncover putative dysfunctions. To date, very little is known on the role of DC subsets in COVID-19 ^31^. A scRNAseq atlas study of PBMC in severe COVID-19 patients identified inflammatory monocytes defective for MHC II molecules ^22^, as was previously shown in severe sepsis patients ^32,33^, but was not tailored to provide sufficient resolution into the DC compartments. The challenge is even greater knowing that some DC subsets, such as pDC and CD141(CLEC9A)^+^ DC were depleted from the blood in severe COVID-19 ^34^ . Through dedicated enrichment steps performed immediately after blood sampling, we were able to capture sufficient cell numbers to define molecular profiles and identify specific defects in all known DC subsets.

As most immune cells, DC are not limited to a single function ^16^. They play a key role in the first line of immune defense by sensing microbial pathogens, and also contribute to direct pathogen control, through the production of anti-microbial peptides ^35^and anti-viral effector molecules ^36^. Other effector functions include the secretion of pro- and anti-inflammatory cytokines, and cytotoxic molecules ^16^. Last, they function as APC to T cells, with which they communicate through a large number of secreted and surface molecules expressed within the immune synapse ^37^. By using scRNAseq, and a combination of supervised and unsupervised bioinformatics methods, we were able to uncover defects in almost all of these processes, in specific APC subsets, associated to COVID-19 severity. This provides the first detailed molecular map of DC subsets and underlying molecular pathways in COVID-19.

Several studies have shown an increase of inflammatory cytokines in severe COVID-19, which may contribute to the severity of the disease ^31^ . Increased circulating levels of IL-1*β* and IL-6 were detected in severe patients ^8,31,38^ . However, the cellular source does not seem to be from circulating cells, but rather from inflammatory monocytes attracted to the lung ^39^, as well as endothelial cells ^10,11^. Our study corroborates these findings for IL-6, with no significant expression detected across APC subsets. However, we did find increased expression of IL-1*β*, CXCL8, and CXCL2, in APC at the global level, and this may contribute to the systemic inflammation. In parallel, we observed an increased TNF signaling in pDC, but decreased in monocytes, suggesting that distinct APC may respond differently to circulating inflammatory mediators.

Type I and III IFN are critical anti-viral cytokines ^21,40^. APC are a central source of IFN following viral sensing. Studies have shown that type I IFN responses are impaired in severe COVID-19 ^8^, which may contribute to persistent viral load. Our data support these findings, since we did not detect any expression of IFN-*α* and IFN-*λ*1 across all APC subsets. However, we were also able to detect critical defects in the response to type I IFN. First, the expression of IFNAR1 and 2 were globally decreased in APC subsets from severe patients. Second, most downstream ISG (both anti-viral, and regulators of IFN signaling) were expressed at lower levels in severe patients, as compared to healthy controls, which themselves are expected to express low levels of ISG given the absence of innate stimulation. Overall, the IFN pathway is defective in severe COVID-19 APC at several levels: IFN production, receptor expression, and downstream ISG responses.

PDC represent a cell type that is highly specialized in anti-viral immunity by producing large amounts of all type I IFN ^41^. Circulating pDC were shown to be diminished in COVID-19 ^34^, but the underlying mechanisms remain unknown. We identified increased expression of pro-apoptotic molecules in pDC from severe COVID-19 patients, associated with decreased relative proportions of pDC. This suggests that pDC could be globally altered through increased cell death. In a separate study, we have shown that *in vitro* SARS-CoV-2 stimulation of normal pDC leads to improved cell survival ^42^, suggesting that the increased apoptosis we observed in severe COVID-19 patient pDC was not due to direct virus-induced killing. In parallel, we detected several defects in various pDC functions: decreased innate sensing, through loss of TLR-7 and DHX36, which are key viral sensors ^43,44^, decreased anti-viral effector functions and cytotoxicity. Hence, we report multi-process defects affecting key aspects of pDC anti-viral functions.

Transcriptomic data, including scRNAseq, allows for the application of methods to infer TF activity, as a way to provide potential upstream mechanisms. We found that several important TF activities were decreased in CD1^+^ DC, suggesting defective immune effector functions in severe COVID-19 patients. STAT2 activity downregulation may indicate a deficiency to cross-present to CD8+ T cells and license their cytotoxic function ^45^. Subversion of DC immunogenicity by targeting STAT2 was observed in other viral infections. ZIKV evades type I IFN responses by antagonizing STAT2 phosphorylation ^46^. Low estimated activity of ESR1, CIITA, USF1, and RFX5, in CD1^+^ DC, may coordinately explain the decrease in MHC II molecules we observed in severe COVID-19 patients, through decreased trans-activation of MHC II promoter ^29,47^. Last, the low activity of EGR1 and RUNX1 TF in severe patients CD1^+^ DC may contribute to an impaired function in CD8 T cell activation and induction of IFN-*γ* ^48,49^. Collectively, our results suggest that several aspects of CD1^+^ DC effector functions may be altered through decreased activity of key TF controlling MHC II expression and T cell stimulation.

Our study provides a unique insight into the physiopathology of APC in severe COVID-19, uncovering previously unknown defects in multiple functional pathways, related to both innate and adaptive immunity. We were able to map molecular pathways in rare DC subsets, many of them previously unexplored in the context of COVID-19 patients. Combined with studies in other anatomical sites ^31^, in particular the lung ^39^, and other disease severity stages, our results should contribute to a better understanding of COVID-19 immunopathology. They also open interesting perspectives for clinical applications. Simple molecular markers of defective APC subsets may be considered as prognostic and stratification biomarkers. This hypothesis echoes with the immune pathology of bacterial sepsis in which multiple defects in APC have already been described ^32,50^. Persistent decrease in circulating dendritic cells, as well as monocyte deactivation as assessed by decreased HLA-DR expression or decreased CD74 mRNA expression, are already known to be predictive of ICU-acquired superinfections in patients with bacterial sepsis ^51–53^. It would be interesting to explore whether such markers, for example pDC apoptosis or CD1^+^ DC MHC II downregulation, appear earlier in the course of COVID-19 and may predict aggravation. From a therapeutic standpoint, many innate adjuvants have been developed to target multiple or specific DC subsets ^54,55^, and could be considered as personalized immunotherapies depending on patient-specific DC dysfunction.

Last, DC are being considered in preventive vaccine development (ClinicalTrials.gov: NCT04386252). Ultimately, our study may form the ground for novel therapies to restore defective APC functions in COVID-19 patients.

## Material and methods

### Patient characteristics and recruitment into the study

This study was part of the Dendrisepsis project aimed at investigating the functional profiles of APC in patients with sepsis. The study was approved by the appropriate institutional review board and independent ethics committee (CPP #2018-A01934-51). We included adult healthy controls, and patients with PCR-proven COVID-19 pneumonia, within 48 hours of admission to intensive care units (ICU) or to the pulmonology department from an urban tertiary care center. Exclusion criteria were the following: haematological malignancy or significant history of bone marrow disease, HIV infection, any immunosuppressive drugs, bone marrow or solid organ transplant recipients, leucopenia (<1000/mm^3^) except if due to COVID-19, pregnancy. With respect to healthy controls, exclusion criteria were the following: history of inflammatory disease, corticosteroid treatment at any dose, infection symptoms within the previous month. Informed consent was obtained from patients or next-of-kin. Patients were classified into moderate pneumonia if requiring oxygen supply < 10 L/min, and severe pneumonia if requiring invasive mechanical ventilation or oxygen supply ≥ 10 L/min. Patients were sampled at admission (day 1) at day 4. Healthy controls were sampled once. Detailed patient characteristics are provided in Table 1 and Supplementary Table 1.

### Cell purification

Blood samples (20ml) were collected from each patient at days 1 and 4 post-hospital admission, and from HC. PBMC were isolated by centrifugation on a density gradient (Lymphoprep, Proteogenix). After FICOLL (GE Healthcare and Lymphoprep StemCell) gradient centrifugation, total PBMC were enriched in CD14^+^ monocytes using the human CD14 microbeads (Miltenyi Biotec) for positive magnetic selection according to the manufacturer’s instructions. The negative fraction remaining after the positive selection of CD14^+^ cells was used for pan-DC enrichment employing the EasySep human pan-DC enrichment kit (StemCell Technologies). Total pan-DC were resuspended with 20,000 CD14^+^ cells and sent for sequencing. Monocyte and pan-DC enrichment were performed immediately after sampling. To avoid DC-T cell clusters that often form in DC-enriched preparations, EDTA-containing medium (DPBS 1X, 0.5% EDTA, 1% Human Serum) was used for sample enrichment in the 4 first patients and one HC (n=8 samples, discovery set, Figures 1-8). Because EDTA can decrease Reverse Transcription efficiency, we validated the findings by using RPMI for all enrichment steps (1640 + Glutamax, 2% BSA, 1% Penicillin/Streptomycin, 1% Sodium Puryvate, 1% MEM NEAA) in the remaining 3 patients and one HC (n=6 samples, validation set, Supplementary Figures 1-5).

### Isolation and Preparation of single cell suspensions

Cell suspensions were subjected to in Gel Bead Emulsion using Chromium 10x Genomics controller according to manufacturer guidelines. To perform single cell RNA sequencing after cDNA amplification, the concentration of each sample was measured using Tapestation 2200 (Agilent). To prepare the cDNA libraries for 10x Genomics Chromium controller, we used the single-cell 3′ v3.1 kit. QC libraries were performed using Tapestation 2200 (Agilent). Illumina Novaseq6000 (100cycles cartridge) with a sequencing depth of at least 50,000 reads per cell was used for sequencing. The input number of cells was estimated at 20,000 cells/ samples.

### Quality control and pre-processing of expression matrices

The raw scRNA-seq fastq files were processed using CellRanger 3.1.0 from 10X Genomics technology and aligned to Grch38 reference genome. Bam files and filtered expression matrices were generated using “cellranger_count”. All expression matrices were loaded into R 4.0.0 using “Read10X” function from Seurat library (https://github.com/satijalab/seurat) version 3.1.5. The latter library was used to perform the analysis workflow.

Pre-processing steps were applied to remove genes expressed in less than 20 cells, and remove cells with fewer than 50 genes or displaying more than 50% mitochondrial transcripts. To minimize technical confounding factors related to the sequencing steps, we evaluated the violin plot distribution of the number of Unique Molecular Identifiers (nUMI), along with the total number of detected genes (nFeatures) per cell for all samples. two upper cutoffs of 6,000 and 50,000 were manually set for the nUMi and nFeatures respectively for each sample. These quality-control metrics filtered-out low quality cells. Normalization to 10,000 reads, centering, and scaling were sequentially applied on the expression matrices to correct for the sequencing depth variability. To reduce computational time of samples integration, we filtered-out cells from other cell types than APC. Cell type annotation is detailed in (Part3). To decipher specific alterations occurring in each specific APC subset, we separately sub-clustered each cell subtype, scaled the data and applied graph-based clustering to obtain cell clusters. Genes encoding for immunoglobulins were removed prior to performing the sub-clustering step for each cell type to get rid of ambient RNA.

### Integration of individual cell matrices into a merged expression matrix from all the samples

To allow comparison across severity states, we integrated the whole expression matrices from all the samples using SCTransform algorithm. Integration anchors, retrieved from the first 20 principal components using the “FindIntegrationAnchors” Seurat function, were then used to integrate the datasets using “IntegrateData()” function. This crucial step added an “integrated” assay to the Seurat Object. Scaling and Principal Component Analysis (PCA) dimension reduction were performed on the integrated assay with 50 principal components. High resolution (resolution=0.8) graph-based clustering, and Uniform Manifold Approximation and Projection (UMAP) dimension reduction were conducted to retrieve and visualize cell clusters. ICA dimension reduction was specifically performed for CD14^+^ monocytes, using 30 dimensions.

### Manual annotation of cell types

Cells were manually annotated based on their expressing levels of their respective set of cell type markers, defined as “cell-type signatures”. For each cell-type signature, enrichment scores were computed using “AddModuleScore()” function per cell with 100 randomly selected control genes, split on 25 bins. Each cell cluster was annotated with a particular cell type if its signature-score median value > 0. Cell-type signatures included: pDC: expression of (“TCF4”, “CLEC4C”, “IRF7”, “IRF8”, “LILRA4”, “IL3RA”, “TLR9”, “SPIB”), cDC (“ANPEP”, “CD1C”, “ITGAX”, “CST3”, “FCER1A”), Monocytes(“CD14”, “FCGR1A”,“ S100A12”, “FCGR3A”, “MS4A7”, “LYZ”, “CXCR3”), AS-DC (“AXL”, “SIGLEC6”,“CD22”), NK cells (“NCAM1”, “FCGR3A”, “GNLY”, “XCL1”, “XCL2”, “NCR1”, “NKG7”), T cells (“CD3D”, “CD3E”, “CD3G”), B cells (CD19”, “MS4A1”, “CD79A”, “CD79B”), Plasma cells (“IGHG2”, “IGHG1”, “IGLC2”, “IGHA1”, “IGHA2”, “IGHA3”, “JCHAIN”, “IGHM”, “XBP1”, “MZB1”, “CD38”, “IGLL5”), Erythrocytes c(“HBB”, “HBA1”), Platelets (PPBP). Cells that were annotated as non-APC were discarded for each sample, before integration, to avoid high computational load during integration step. For Monocytes and cDCs, a subsequent classification of cells was performed according to their expression levels of monocytes and cDC subset markers (CD14, FCGR3A for monocytes, CD1C and CLEC9A for cDCs).

### Statistical analysis

Differential expression analysis between severity groups was performed using the “FindAllMarkers()”Seurat function, using Wilcoxon-ranked test and a cutoff was set to logFC>0.3 to filter out false positive differentially expressed genes. P-values were corrected using Benferroni correction method. Only tested are the genes which are detected in a minimum fraction of 20% of each severity group. Median values of violin plot distributions of either gene expression levels or pathway-enrichment scores were compared using wilcoxon test, taking as a reference the Healthy Control (HC). Note that the statistical calculations for the violin plot distribution is derived from the cell count in expression values / enrichment scores comparisons.

### Pathway enrichment analysis

Pathway enrichment analysis was performed to seek for the perturbed or enriched pathways in severity groups, as compared to the HC. Human MsigDB hallmark signatures (https://www.gsea-msigdb.org/gsea/msigdb/index.jsp) were loaded into R session using the “msigdbr” library version 7.0.1, the category was set to “H” for “Human”. Enrichment test was performed using the “enricher” function, from the “gprofiler” version 3.16.0. Msig Database hallmark signatures were given as input to the “enricher” function. P-values were corrected using Benferroni correction method. Encoding genes for each enriched pathway were extracted and used as module to construct a “pathway-score” signature using “AddModuleScore” from Seurat library.

### Transcription Factor activity inference

We sought to decipher variation of transcription factor (TF) activity between severity groups within particular cell types to avoid capturing differentially active TF related to lineage markers. Dorothea (https://saezlab.github.io/dorothea/) ressource was used to infer TF activity. In this context of single cell level resolution, we constructed regulons based on the mRNA expression levels of each TF from a manually curated database, along with the expression level of its direct targets. We used Hence, TF activity is set to be a proxy of a transcriptional state of its targets. We created TF regulons using “dorothea_regulon_human” wrapper function from “dorothea” library version 0.99.10, and chose “A” and “B” high confidence TF selection. Viper scores were computed on the dorothea regulons, scaled and added as “Dorothea” slot on the integrated Seurat object. To allow comparison of TF score activities, mean and standard deviation values of the scaled viper scores were computed per severity group. Transcription factors were ranked according the variance of their corresponding viper scores. Top 50 highly variable score scores per severity groupe (n=150 TF in total) were kept for visualization of their corresponding scores.

### Manual construction of functional signatures

To evaluate the dysregulations occuring at the functional level for each APC subset from COVID-19 patients, we established a manually curated list of effector genes involved in specific APC functions, including: “Attraction”, “Anti-viral effector molecules”, “Cytotoxicity”. The signature construction relied on a thorough mining of existence literature, using a combination of MeSH terms and keywords on the PubMed search tool. Each selected molecule was considered as “effector” of the related function if there is at least one experimental proof in a human setting. Overall, we outlined 12 “Cytotoxicity” effector molecules, 29 “Anti-viral” effector molecules along with 18 “Attraction” effector molecules.

“Innate sensing” effectors included 12 genes (DDX58, DHX58, CGAS,IFI16,AIM2,IRF3, TMEM173,NLRP3,PYCARD,TLR7,TLR9,DHX9,DHX36), and were from ^56,57^ . Both “regulators of Interferon signalling” and “Anti-viral ISG” were implemented through a literature mining of a pioneer article, from ^25^.

### Analysis of intercellular communication networks

Communication scores were generated using ICELLNET R package (https://github.com/soumelis-lab/ICELLNET/master). This package allows computation of cell-cell communication scores between cell subsets, given their corresponding transcriptomic profiles from the same of different datasets. Considering severity groups separately, only clusters including more than 15 cells were considered for the analysis. The average gene expression profiles of APC subset clusters were given as input to ICELLNET package, to compute communication scores between APC subsets and T lymphocytes for each severity group. As our datasets did not include T cells for the analysis, we used as reference the T lymphocyte transcriptomic profile from the Human Primary Cell Altas included in ICELLNET package (n=39). From ICELLNET ligand-receptor interaction database, we only selected the 144 interactions known to be involved in DC-T communication^58^ . Barplot and dotplot representations were generated to compare the proportions of communication type scores (checkpoint, cytokines, chemokines) between severity groups.

## Acknowledgments

We thank our patients for participating into this study. We also wanted to acknowledge Dr Nathalie Marin, Dr Mariana Cojocaru, Dr Nicolas Carlier, Dr Jonathan Marey, Prof. Dominique Monnet and Dr Tali-Anne Szwebel (all from Cochin hospital) for their help in the inclusion of patients and controls. We thank A. Mohammed for performing the sequencing runs on cellRanger at the sequencing platform of Institut du Cerveau et de la Moelle épinière (ICM). We thank Dr. Martine Blot, Dr. Florence Niedergang and Dr. Evelyne Lauret from the Institut Cochin for their precious help during the setting up of the study. The COVID-19 emergency plan from Université de Paris supported this study in technical aspects regarding the efficient processing of sequencing the samples. This study was supported by the ANR DENDRISEPSIS (ANR-17-CE15-0003), ANR APCOD (ANR-17-CE15-0003-01), and Université de Paris « PLAN D’URGENCE COVID19» grants. L.M was supported by a PhD fellowship from La Ligue Contre le Cancer, and E.A by a PhD fellowship from Servier.

## Notes

### Competing Interest Statement

The authors have declared no competing interest.

