## Supplementary files and Table for "Single cell RNA sequencing of blood antigen-presenting cells in severe Covid-19 reveals multi-process defects in antiviral immunity"

### Supplementary Figure 1

**a**

| Sample name | Time | Severity state | CD14+ Monocytes | pDC | CD1c+DC | CD16+ Monocytes | CLEC9A+ DC | AS-DC |
|---|---|---|---|---|---|---|---|---|
| Healthy_C | / | Healthy | 1936 | 2287 | 1446 | 383 | 201 | 109 |
| j0301.J1 | D1 | Severe | 210 | 20 | 11 | 11 | 0 | 1 |
| j0301.J4 | D4 | Severe | 2232 | 637 | 1068 | 368 | 37 | 20 |
| j037.J1 | D1 | Severe | 423 | 191 | 242 | 21 | 8 | 19 |
| j037.J4 | D4 | Severe | 437 | 107 | 240 | 12 | 5 | 15 |
| j0501.J1 | D1 | Moderate | 3817 | 275 | 445 | 276 | 23 | 35 |
| j0501.J4 | D4 | Moderate | 2443 | 415 | 321 | 273 | 25 | 14 |
| j0502 | D1 | Moderate | 307 | 26 | 12 | 10 | 2 | 0 |
| Healthy_C2 | / | Healthy | 1437 | 6 | 520 | 90 | 39 | 166 |
| j0142.J1 | D1 | Severe | 1962 | 2 | 43 | 30 | 2 | 16 |
| j0142.J4 | D4 | Severe | 543 | 0 | 1 | 22 | 0 | 3 |
| j0143 | D1 | Severe | 361 | 14 | 24 | 144 | 2 | 16 |
| j0401.J1 | D1 | Mild | 84 | 128 | 167 | 2390 | 5 | 7 |
| j0401.J4 | D4 | Mild | 14 | 29 | 151 | 1587 | 4 | 88 |

**b**

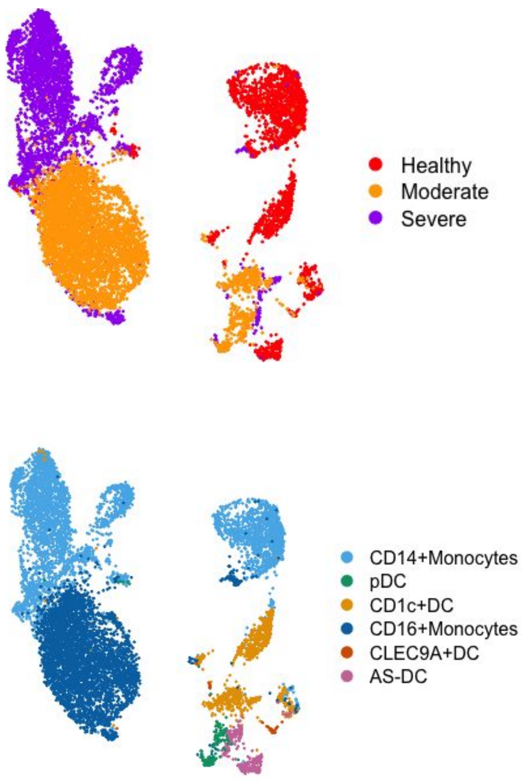


**c**

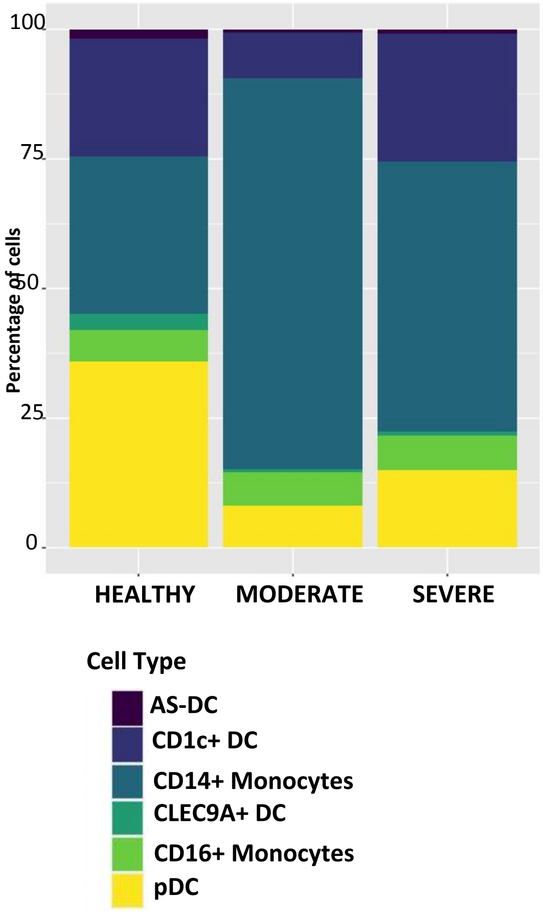

**Supplementary Figure 2**

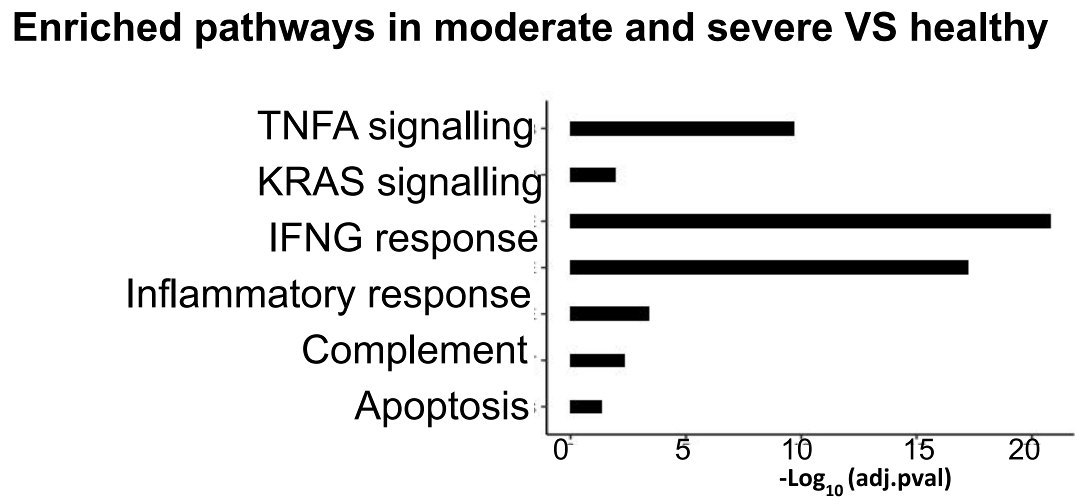


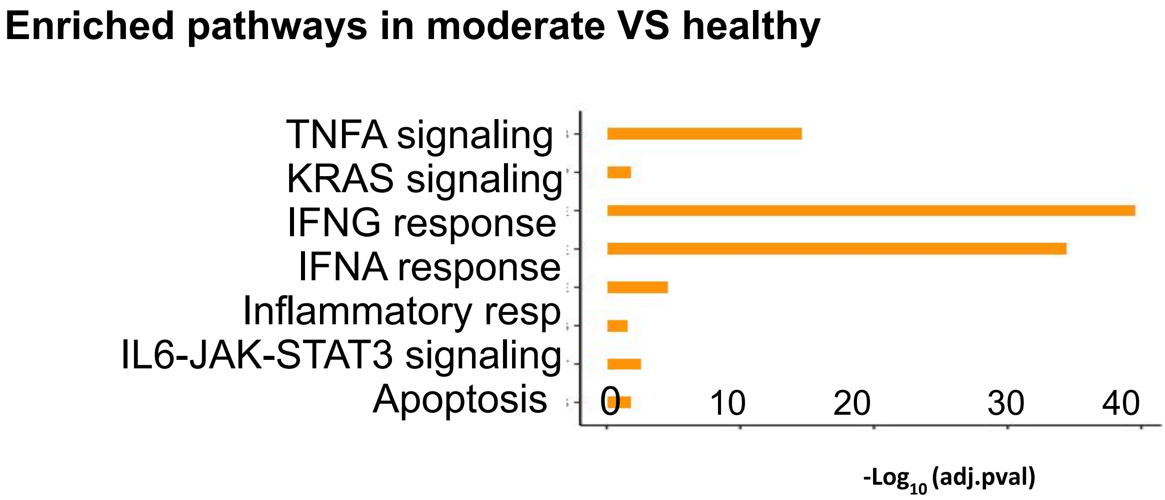


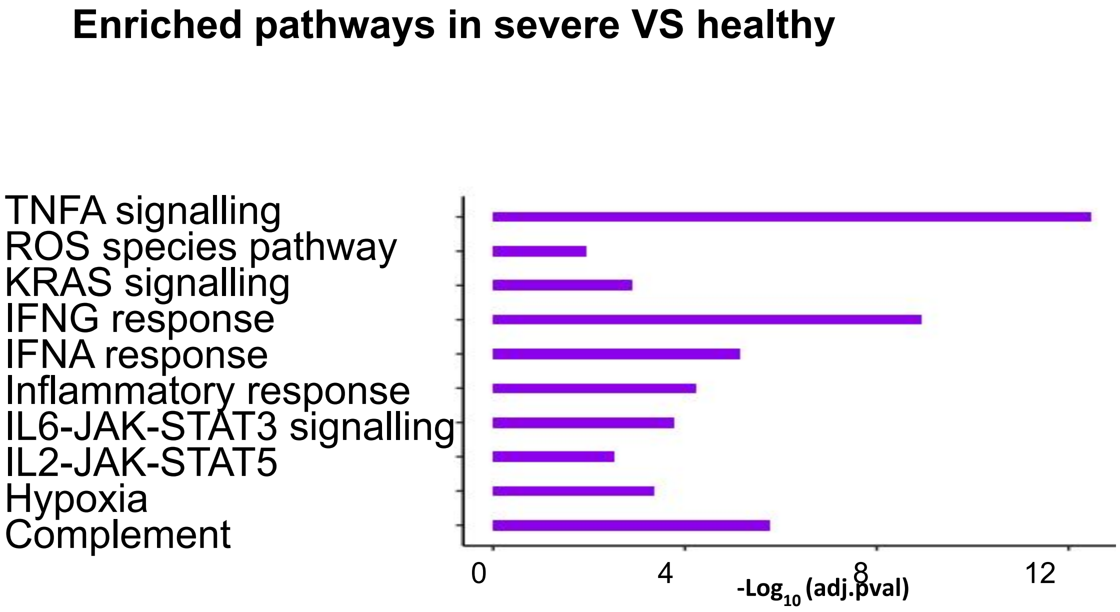

Supplementary Figure 3

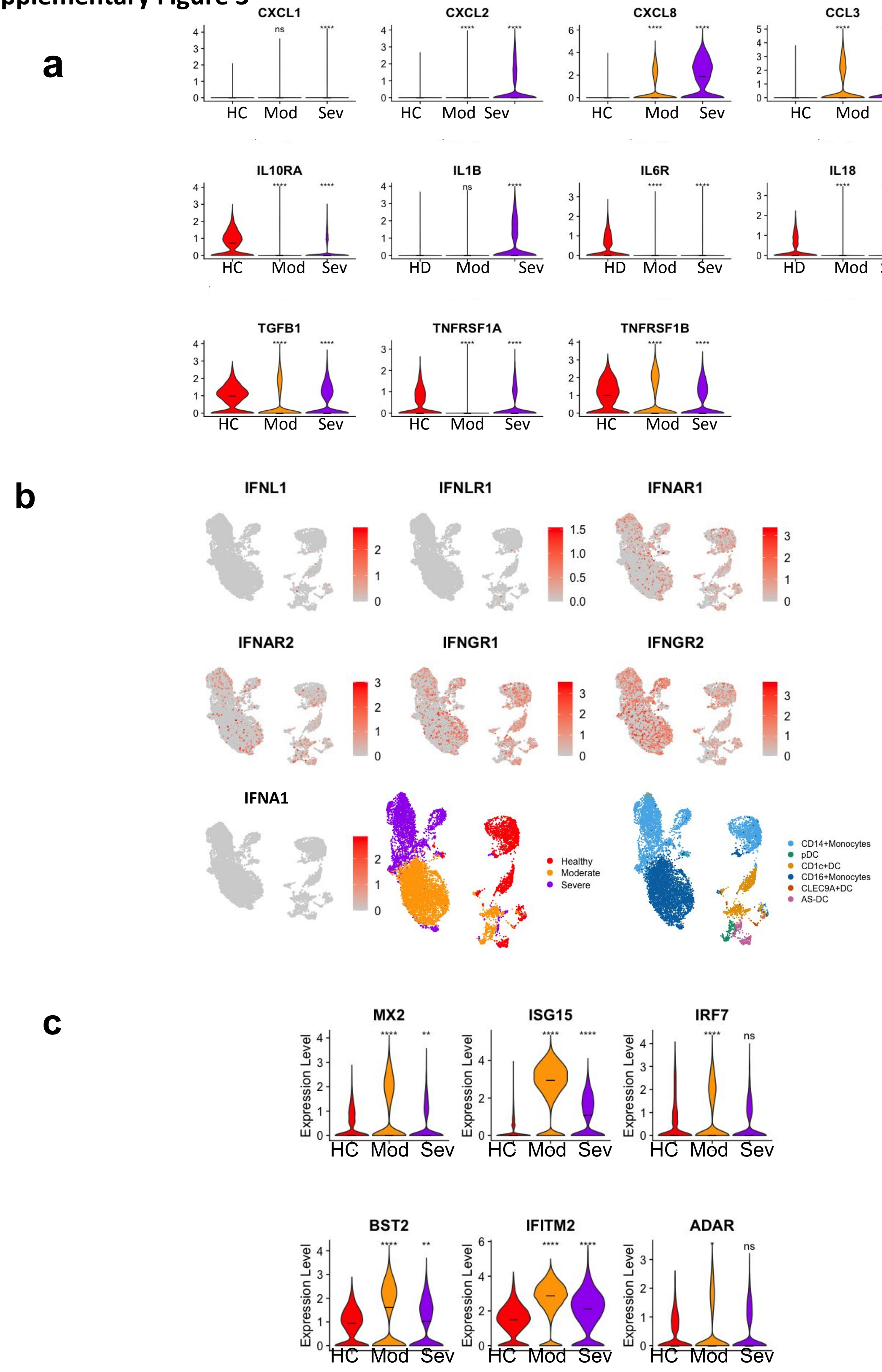

**Supplementary Figure 4**

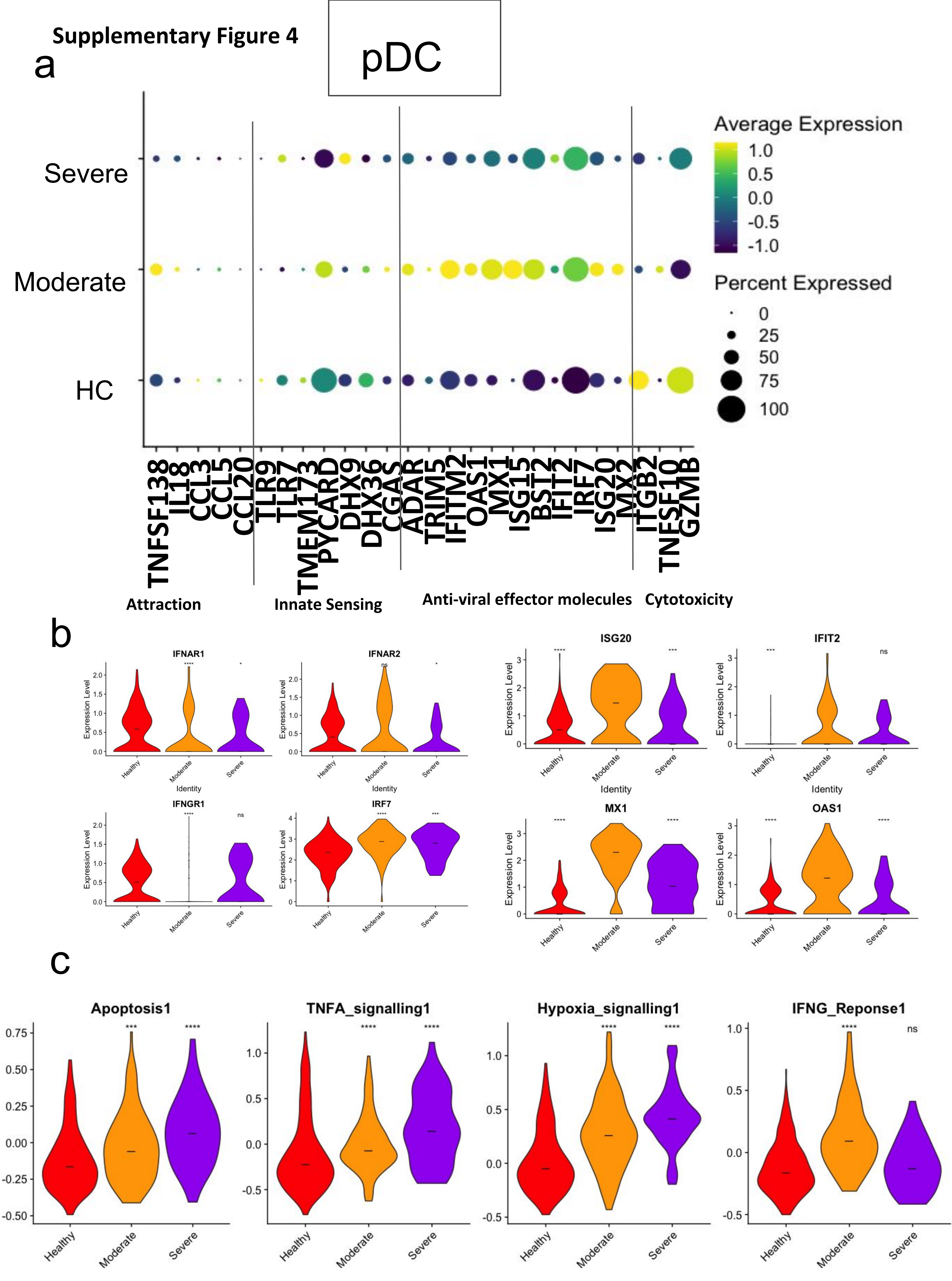

Supplementary Figure 5

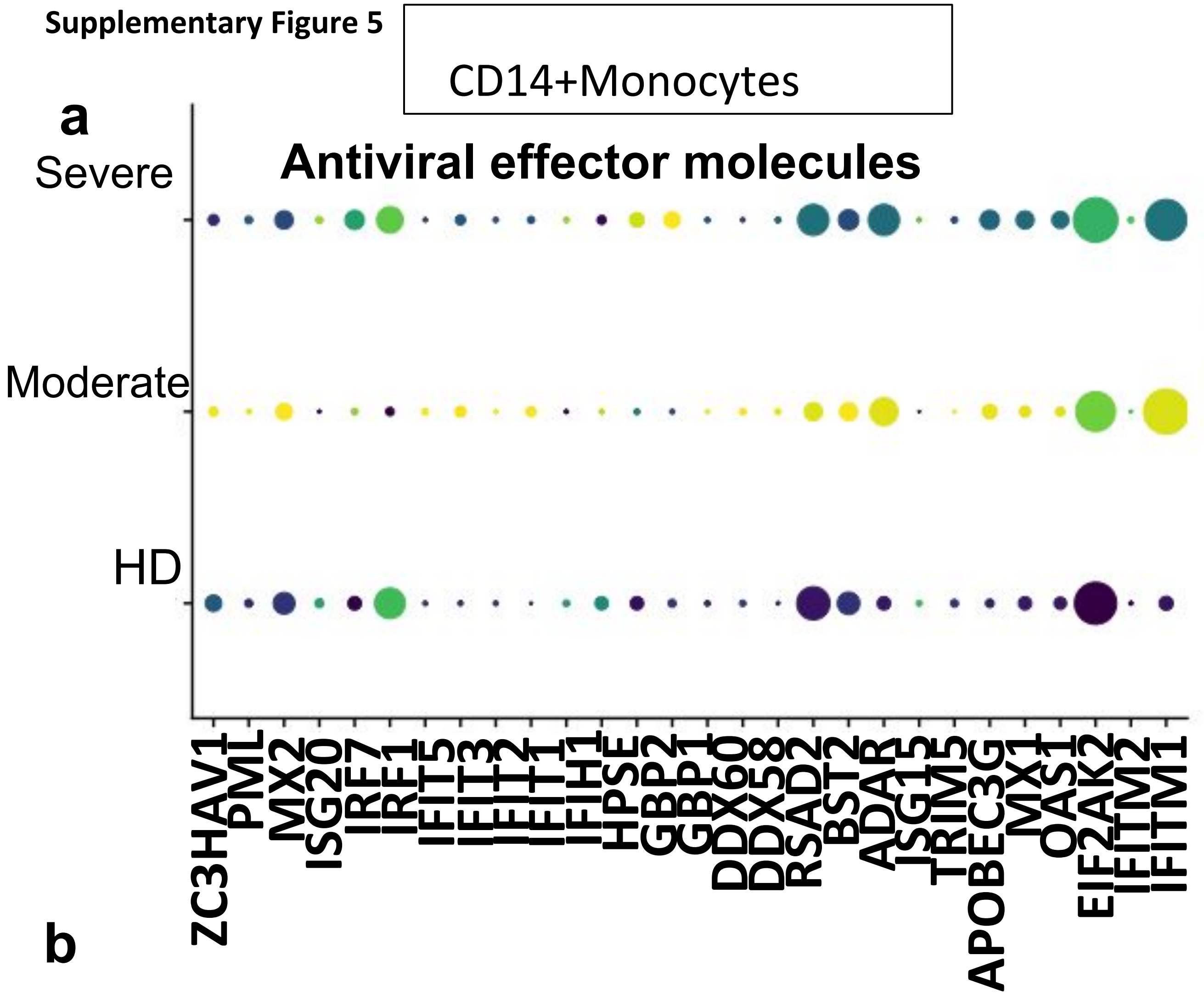


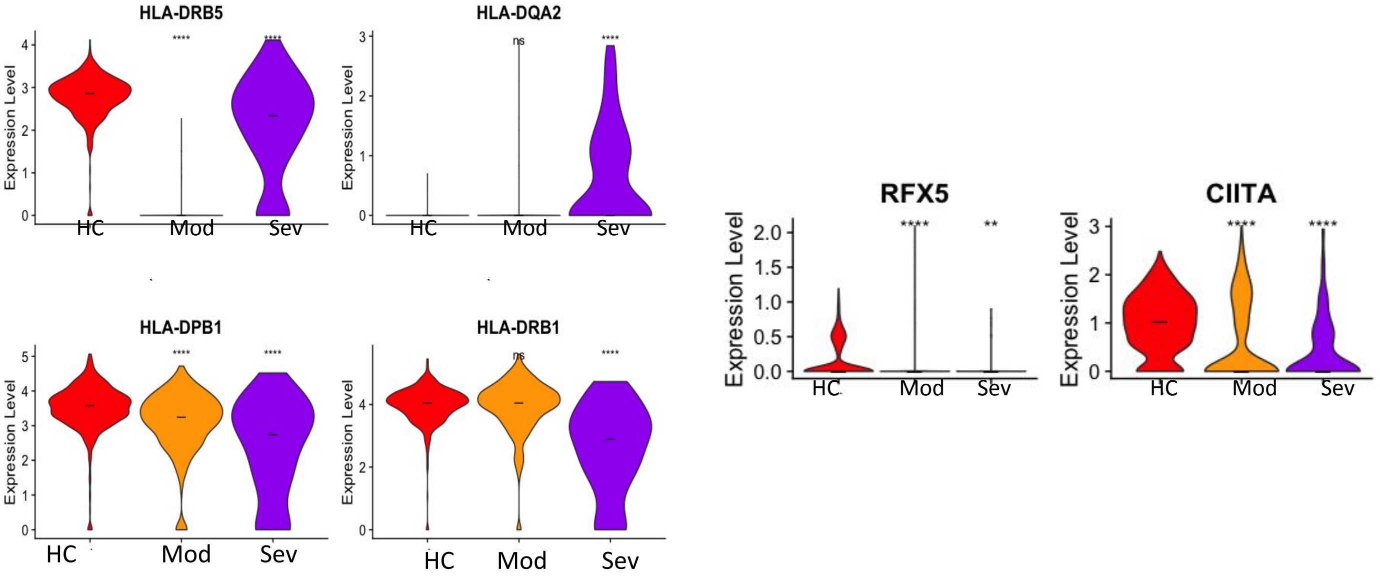

| Subjects | Gender | Age (years) | Pneumonia severity | Baseline sample | | | | | | | | | Day 4/5 sample | | | | | | | | | Outcome |
|---|---|---|---|---|---|---|---|---|---|---|---|---|---|---|---|---|---|---|---|---|---|---|
| | | | | Ventilatory support | Concurrent bacterial infection | Leucocytes (G/L) | Neutrophils (G/L) | Monocytes (G/L) | Lymphocytes (G/L) | Platelets (G/L) | CRP (mg/L) | PCT (µg/L) | Ventilatory support | Concurrent bacterial infection | Leucocytes (G/L) | Neutrophils (G/L) | Monocytes (G/L) | Lymphocytes (G/L) | Platelets (G/L) | CRP (mg/L) | PCT (µg/L) | |
| **Patient #1** | Male | 73 | Severe | Invasive mechanical ventilation | No | 8.9 | 6.56 | 0.3 | 1.53 | 226 | 157 | 0.84 | Invasive mechanical ventilation | no | 6 | 4.68 | 0.68 | 1.3 | 161 | 95 | 0.64 | Survived |
| **Patient #2** | Male | 53 | Severe | Invasive mechanical ventilation | No | 4.45 | 3.4 | 3.1 | 0.6 | 246 | 208 | 0.69 | Invasive mechanical ventilation | no | 7.5 | 6.33 | 0.35 | 0.63 | 346 | 36.7 | ND | Survived |
| **Patient #3** | Male | 85 | Mild/moderate | O2 flow6 L/min | No | 9.75 | 7.71 | 1.07 | 0.81 | 257 | 230 | 0.13 | O2 flow6 L/min | yes (UTI Citrobacter) | 7.59 | 5.86 | 0.83 | 0.78 | 309 | 127 | 0.11 | Survived |
| **Patient #4** | Female | 88 | Mild/moderate | O2 flow3 L/min | No | 3.4 | 2.4 | 0.14 | 0.78 | 276 | 32 | 0.08 | | | | | | | | | | Survived |
| **Patient #5** | Male | 66 | Severe | Invasive mechanical ventilation | No | 6.4 | 5.43 | 0.21 | 0.6 | 227 | 266 | 0.27 | Invasive mechanical ventilation | yes (VAP MSSA) | 7.6 | 6.66 | 0.46 | 0.49 | 252 | 260 | 6.62 | Survived |
| **Patient #6** | Male | 94 | Severe | O2 flow15 L/min | No | 6.48 | 5.35 | 0.26 | 0.8 | 224 | 169 | 0.07 | | | | | | | | | | Died |
| **Patient #7** | Male | 66 | Mild/moderate | O2 flow2 L/min | No | 7 | 4.89 | 1 | 0.94 | 255 | 113 | 0.08 | Room air | no | 8.43 | 5.52 | 1.1 | 1.75 | 419 | 12.4 | 0.08 | Survived |
| **Healthy control #1** | Male | 46 | | | | | | | | | | | | | | | | | | | | |
| **Healthy control #2** | Male | 73 | | | | | | | | | | | | | | | | | | | | |

Supplementary table : Individual characteristics of seven patients and two healthy controls
Patient #4 refused to be sampled at day 4/5. Patient #6 was not sampled at day 4/5 because in palliative care.

Abbreviations : UTI, urib=nary tract infection ; VAP, ventilator-associated pneumonia

### Supplementary Figure Legends

**Supplementary Figure 1. Cellular map of APC subsets constructed from COVID-19 patients**

**a.** Cell counting of different APC populations identified by single-cell RNA sequencing analysis in all samples from the discovery and validation sets, **b.** Cellular map of APC subsets at the single-cell resolution level from the validation set based on either severity or APC subsets, **c.** Proportions of APC subsets within severity groups from the discovery set; cell type subsets are color-coded.

**Supplementary Figure 2. Increase of inflammation related pathways in APC subsets from COVID-19 patients**

Pathway enrichment analyses performed on Differentially Expressed Genes (DEG) between severity groups in the validation set. P-values were adjusted to multiple tests using Bonferroni correction method.

**Supplementary Figure 3. Perturbation of cytokine and IFN response levels in severe COVID-19 APC from the validation set**

**a.** Violin plot representation of gene expression levels for cytokines (IL1B, TGFB1, IL-6 and IL-18), chemokines ( CXCL1,CCL3, CXCL8 and CXCL2) and receptors (IL6R, IL10R and TNFRSF1A) detected in the validation set, and comparison of their corresponding distributions in severe and moderate patients to the Healthy Control (HC), **b.** Feature plot of IFNs and their receptor expression across all cell types and severity cases within UMAP (discovery dataset). Expression levels are color-coded, **c.** Violin plot representation of anti-viral IFN stimulating genes between severity groups. Asterisks above severe indicate *P* values for severe versus control; asterisks above moderate indicate significance of moderate versus control. $*P < 0.05$, $**P < 0.01$, $***P < 0.001$.

**Supplementary Figure 4. Global defects in pDC-related functions in severe patients correlated with increased apoptosis**

**a.** Dot plots of pDC-related functions “Attraction”,”Innate sensing”, “Anti-viral effector molecules”, “Cytotoxicity” in pDC from HC, moderate and severe patients. Expression levels are color-coded; Percentage of cells expressing the respective gene is size coded, **b.** Violin plot representation of gene expression for IFN receptors (IFNAR1 and 2), IRF7, and anti-viral effector molecules, **c.** Violin plot representation of upregulated pathways in COVID-19 patients, and comparison between the three severity groups. Asterisks above severe indicate *P* values for severe versus control; asterisks above moderate indicate significance of moderate versus control. Exceptionally, for ISG20, IFIT2, MX1, OAS1, asterisks above severe indicate *P* values for severe versus moderate; asterisks above HC indicate significance of HC versus moderate. $*P < 0.05$, $**P < 0.01$, $***P < 0.001$.

**Supplementary Figure 5. Anti-viral properties along with MHC-II antigen presentation are defective in CD14+ monocytes and CD1c+DC respectively**

**a.** Dot plots of “anti-viral effector molecules” in CD14+ monocytes from HC, moderate and severe patients. Expression levels are color-coded; Percentage of cells expressing the respective gene is size coded, **b.** Violin plot representation of class II antigen presentation genes in CD1c+DC from COVID-19 patients and the HC from the validation set, and comparison between the three severity groups. Asterisks above severe indicate *P* values for severe versus control; asterisks above moderate indicate significance of moderate versus control. $*P < 0.05$, $**P < 0.01$, $***P < 0.001$.

**Supplementary Figure 6. Perturbation of DC-T communication in severe COVID-19 patients from the discovery set.**

**a.** Connectivity maps describing outward communication from APC from single cell dataset at day 1 to T lymphocytes (n=39), according to patient severity (healthy, moderate, severe). T lymphocytes transcriptomic profiles are from Human Primary Cell Atlas, included ICELLNET R package. For APCs, average cluster gene expression profiles were considered. Only DC-T interactions are taken into account to compute communication score (manually curated, n=144 interactions), **b.** Barplot of each communication score with contribution by families of communication molecules for outward communication from APCs to T lymphocytes, **c.** Focus on CD1c+ DC outward communication to T lymphocytes, representing specific individual interaction scores that differ from at least 10 between moderate and severe patients (cutoff chosen for clarity purpose).

**Supplementary Table legend**

**Table 1. Individual characteristics of seven patients and two healthy controls included in our study**
